# Fluorescence lifetime estimation: a practical approach using Flipper-TR FLIM

**DOI:** 10.1101/2025.09.23.678124

**Authors:** Tithi Mandal, Aurélien Roux, Juan Manuel García-Arcos

## Abstract

Flipper-TR is a membrane dye sensitive to lipid packing widely used to probe membrane tension in live cells via fluorescence lifetime imaging microscopy (FLIM). However, no consensus currently exists on the optimal strategy for extracting lifetime values, particularly across varying experimental setups and biological systems. Here, we systematically compare multiple approaches to estimate Flipper-TR lifetime, including multi-exponential reconvolution fitting, tail fitting, mean photon arrival time (first moment), and phasor analysis. These estimators are tested against changes in photon budget, sample characteristics, microscope manufacturer, and laser frequency. This offers a comprehensive benchmark and decision-making framework for quantitative FLIM analysis of Flipper dyes in various contexts.

## 1. Introduction

### 1.1 What is membrane tension?

Biological membranes are dynamic interfaces that regulate interactions between cells and the environment, serve as platforms for protein scaffolds, or separate intracellular compartments. The main modulator of membrane mechanics is membrane tension, defined as the stress resulting from changing the apparent surface area of the membrane (Helfrich, 1973). Plasma membrane tension plays a central role in processes such as cell migration (Gabella et al., 2014; Hetmanski et al., 2019; Houk et al., 2012; Keren et al., 2008; Kozlov & Mogilner, 2007; Lieber et al., 2013, 2015), collective migration (Chakraborty et al., 2022), and regulates membrane signaling (Bergert et al., 2021; De Belly et al., 2021), actin polymerization (Mueller et al., 2017), axon growth (Dal & Sheetz, 1995; Shi et al., 2022), and cell spreading (Raucher & Sheetz, 2000; Venkova et al., 2022).

Several tools have been developed to probe membrane tension *in vitro* or in living systems. Some techniques measure the mechanical properties of the cell surface, in which tension can be one parameter to fit. For example, by using fluctuation spectroscopy using RICM (Reflection Interference Contrast Microscopy), where interference patterns provide nanometer-scale information on membrane height. The spatiotemporal fluctuations of the membrane position are fitted with a theoretical model in which both bending rigidity and membrane tension determine the fluctuation amplitudes. Because tension suppresses long-wavelength fluctuations, the relative contribution of low-versus high-frequency modes allows membrane tension to be quantitatively extracted. (Betz & Sykes, 2012; Biswas et al., 2019).

The most common method for measuring membrane tension involves measuring the force required to pull membrane tethers from different parts of the cell membrane. Tether force measurements are an invasive technique and are limited to one or a few measurements per cell. Importantly, it cannot be used to assess the tension of the membrane in contact with the substrate, because the membrane needs to be accessed to pull a tube from it. Moreover, tether force includes additional contributions from membrane-cytoskeleton interactions and bilayer bending rigidity (Bozic et al., 1997; Cuvelier et al., 2005; Dai & Sheetz, 1999; Evans et al., 1996; Heinrich et al., 1999; Hochmuth et al., 1996), which is lipid composition-dependent (Sayem Karal et al., 2023). Other methods involve the use of fluorescent probes sensitive to lipid packing, such as Flipper-TR. The fluorescence lifetime of Flipper-TR is sensitive to the physico-chemical environment of the membrane which as allowed to measure membrane tension without the constraints imposed by other techniques. In the next sections, we describe how Flipper-TR works and how we can use it as a tension reporter.

### 1.2 How does Flipper-TR work? Photochemistry of Flipper-TR

Fluorescence lifetime describes the average time a fluorophore spends in its excited state before returning to the ground state. Relaxation can occur either by emitting a photon or by ‘dark’ non-radiative pathways such as internal conversion, intersystem crossing, quenching, or energy transfer. A Perrin–Jabłoński diagram (Figure 1A) summarizes these processes by depicting the ground and excited electronic states and the possible transitions between them. Each decay pathway is characterized by a rate constant (k), and the overall decay rate is the sum of all radiative and non-radiative rates. Consequently, the fluorescence lifetime is given by the inverse of this total rate. This means that when additional relaxation channels are available (e.g., quenching, FRET), the total decay rate increases, and the fluorescence lifetime becomes shorter.

**Figure 1:**
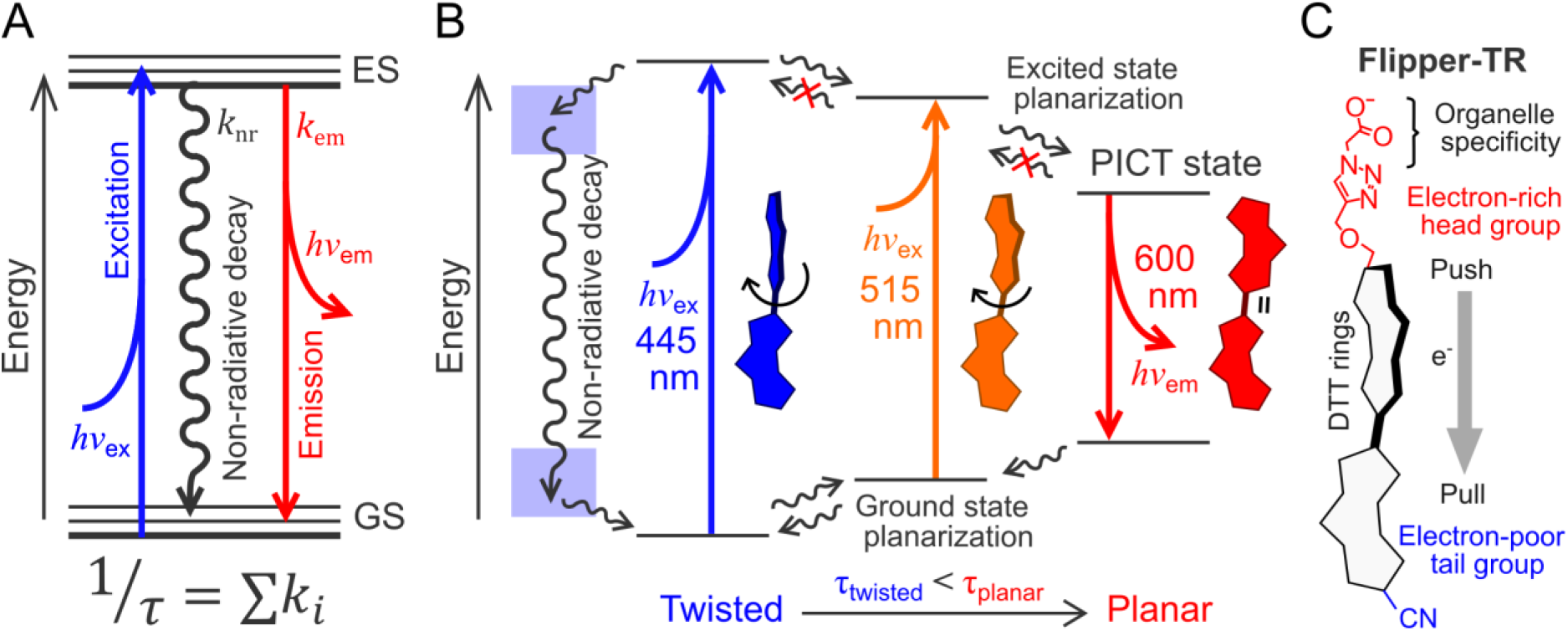
Photochemistry of Flipper-TR. (**A**) Generic Perrin-Jabłoński diagram, describing the electronic and vibrational energy states of a molecule, along with the radiative and non-radiative transitions after light absorption. Energy levels below correspond to the ground state (GS), whereas the energy levels above correspond to the excited state (ES). Molecules transition to excited state levels by absorbing a photon at the excitation wavelength (*hv*_ex_). Molecules can relax and transition back to the GS by non-radiative relaxation (no photon emission) or by emitting a photon at the emission wavelength (*hv*_em_). Each of the decay processes have a rate k [1/ns] involved. The sum of the rates of all decay processes equals the inverse of the fluorescence lifetime. (**B**) Perrin-Jabłoński diagram of flipper probes. Flipper molecules can be planarized at the ground state by external forces and changes in lipid packing. The less planar ground state (blue) has an absorption maximum at 445 nm, whereas the more planar state has an absorption maximum at 515 nm. The more planar excited state undergoes ultrafast excited-state planarization and emission from planar excited state (PICT), which is the only conformation with photon emission. The twisted excited state can also undergo ultrafast planarization and photon emission, but can also lead to ultrafast excited-state deplanarization followed by non-radiative decay through more twisted states. This leads to a lower quantum yield and lower lifetime value for twisted molecules compared to planar molecules. (**C**) Flipper-TR structure-function. The probe’s mechanophore is a push-pull dithienothiophene (DTT) dimer held twisted in the ground state. The red segment denotes the plasma-membrane–targeting headgroup and the donor side of the chromophore (exocyclic donor located adjacent to the headgroup), whereas the blue segment marks the electron-poor acceptor side. In membranes, mechanical packing/compression promotes planarization of the DTT “flippers,” which strengthens donor-acceptor conjugation and the push–pull polarity. The gray arrow (e⁻) indicates the direction of charge displacement upon planarization/ICT from donor (red side) to acceptor (blue side). The headgroup governs localization, while the donor/acceptor elements on the scaffold generate the push–pull that confers mechanosensitivity.

Flippers are small-molecule mechanosensors whose photophysics are coupled to their molecular conformation (Chen et al., 2023). Upon excitation, flippers undergo ultrafast (picosecond) planarization to a planar intramolecular charge-transfer (PICT) state that is the sole emissive state. Planarization competes with other non-radiative (dark) pathways; conditions favoring planarization therefore increase quantum yield and lifetime (Chen et al., 2023). Because absorption can originate from several states, but emission always originates from PICT, pre-planarization red-shifts absorption while leaving the emission maximum nearly unchanged around 600 nm (Figure 1B).

The structure of Flipper-TR, the plasma-membrane-targeting flipper, is tightly linked to its functioning. An anionic headgroup drives plasma-membrane enrichment, conferring a plasma membrane specificity, while other headgroup variants have recently been developed targeting other organelles (Chen et al., 2023). The mechano-sensing core of Flipper-TR consists of two dithienothiophene (DTT) units held in a twisted, non-planar ground state by repulsive chalcogen interactions (Figure 2C). Once inserted in a lipid bilayer, the probe’s twist–planarity equilibrium is influenced by the local physicochemical environment. Membrane conditions that mechanically planarize the molecular scaffold (e.g., tighter lipid packing) enhance conjugation across the push–pull system. As described above, this produces i) a higher quantum yield (more photons emitted), ii) a red shift in absorption/excitation, and iii) lengthens fluorescence lifetime, providing a quantitative readout with fluorescence lifetime imaging microscopy (FLIM). In solvents such as water, the molecule is practically non-fluorescent, eliminating the need to wash the probe during bioimaging.

**Figure 2:**
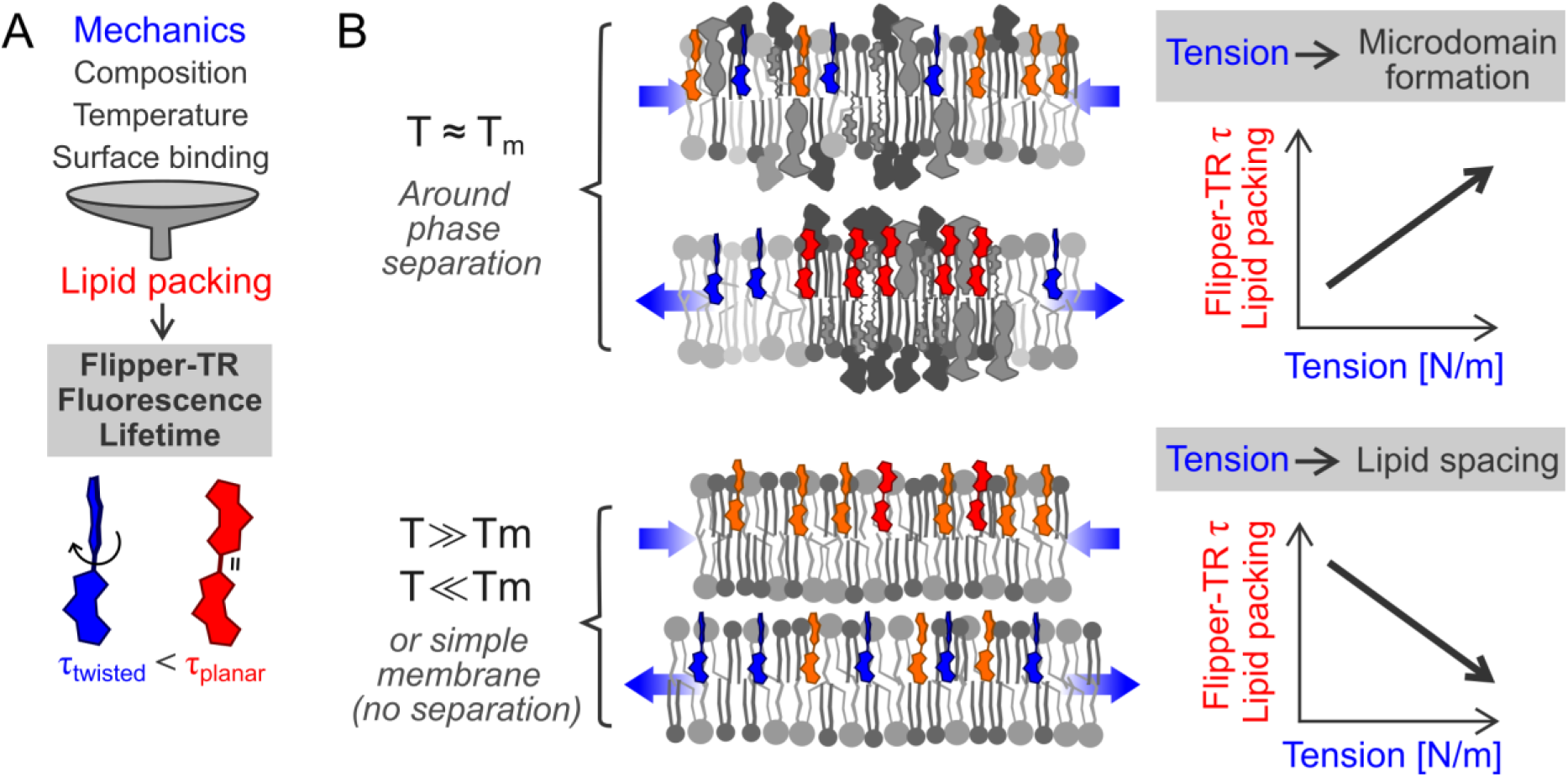
Relationship between membrane mechanics parameters and lipid packing. (**A**) Lipid packing results from a complex interplay between membrane parameters: mechanics, composition, temperature, and other factors such as surface binding. Flipper-TR lifetime directly reports on lipid packing without distinction of the upstream factor dominating changes in lipid packing. The conformation of the probe (blue, twisted; red, planer) results in changes in the fluorescence lifetime (τ), measured by FLIM. (**B**) Effect of tension on lipid packing in different membrane types. In membranes around phase separation, tension can promote microdomain formation. This results in a positive correlation between tension and Flipper-TR lifetime/lipid packing. In membranes far from the transition temperature or in simple membranes not able to phase separate, tension increases space between lipids. This results in a negative correlation between tension and Flipper-TR lifetime/lipid packing.

The probe has been well characterized chemically. Flipper-TR molecules in solution insert and align along the lipid tails in external leaflet of the bilayer (Dal Molin et al., 2015; Licari et al., 2020), without affecting the membrane structure significantly (Neuhaus et al., 2017). Flippers act as monomers and partition almost equally between more and less ordered phases (Dal Molin et al., 2015; García-Calvo et al., 2022). Flipper-TR also show negligible flip-flop (López-Andarias et al., 2021), but can transfer rapidly between different membranes though an aqueous solution (Bayard et al., 2023; López-Andarias et al., 2022), which is useful for staining but important to consider in dynamic systems.

### 1.3 Why do lipid packing probes report membrane tension?

The response of lipid packing probes such as Flipper-TR to membrane tension is intimately linked to an important characteristic of biological membranes: their lateral separation into nanodomains (Eggeling et al., 2009; Honigmann et al., 2014). These nanodomains are characterized by different degrees of lipid packing or lipid order, which refers to the arrangement and density of lipid molecules (Sezgin et al., 2012).

Notably, lipid packing and domain formation are not only linked to lipid composition, phase, and temperature, but also to membrane tension (Figure 2A). This represents an interesting link between the mechanics and the molecular organization of membranes. Increasing membrane tension by hypotonic shocks in model membranes around transition points drives lateral phase separation, promoting the formation of ordered lipid domains and higher lipid packing (Akimov et al., 2007; Hamada et al., 2011; Oglęcka et al., 2014; Rathe et al., 2021; Wongsirojkul et al., 2020; Yanagisawa et al., 2008). Conversely, in membranes far from a demixing transition (i.e., far from the transition temperature *T*_*m*_, *T* ≫ *T*_*m*_ or *T* ≪ *T*_*m*_), or with simple compositions that cannot phase separate, increasing tension leads instead to a decreased lipid packing, as expected from increased lipid spacing (Figure 2B bottom panel).

This effect was measured with Laurdan probes (Boyd & Kamat, 2018; Zhang et al., 2006) or with Flipper-TR (Colom et al., 2018). In cells, results seem consistent with the transition point scenario (*T* ≈ *T*_*m*_, Figure 2B top panel): increasing plasma membrane tension by hypotonic shocks globally promotes ordered domain formation (Ayuyan & Cohen, 2008) and increased lipid packing measured with flipper (Roffay et al., 2021) and Laurdan probes (Zapata-Mercado et al., 2022). Pipette aspiration in cells yielded the same results, correlating as well with tether pulling force (Roffay et al., 2021). Nanodomains have not yet been resolved with Laurdan or flipper probes, but their behavior upon increased membrane tension is consistent with tension-induced domain formation.

The linear correlation between the membrane mechanics and its molecular organization allows for a proxy measurement of membrane tension with lipid packing-sensitive probes such as Flipper-TR, determining the probe conformation by FLIM (Colom et al., 2018; Dal Molin et al., 2015). Importantly, the link of lipid packing to other parameters such as lipid composition, phase, and temperature imposes a careful experimental design and careful controls to understand the main driver behind the measured changes in lipid packing.

### 1.4 Introduction to Flipper-TR lifetime estimators

Fluorescence lifetime imaging microscopy is a technique to distinguish the molecular environment of fluorophores. FLIM is gaining popularity and is used in a broad range of biological problems, from FRET analysis to metabolic imaging. In this section, we focus on FLIM analysis of Flipper-TR. For a broader overview of FLIM principles, methods and analysis please refer to previous publications (Hirvonen & Suhling, 2020; Torrado et al., 2024).

The most common way to perform Flipper-TR FLIM measurements involves the use of a pulsed laser source coupled with a time-correlated single photon counting device (TCSPC). Laser sources usually come at a frequency of 80MHz, so a pulse picker device is used to lower the frequency to 20MHz (50-nanosecond interval), ideal for Flipper-TR measurements. The excitation wavelength is around 488 nm, while the emission band pass is 600/50 nm. It is important to control these parameters, as lifetime is sensitive to the emission wavelength (Ragaller et al., 2024a). TCSPC devices record the time delay between the excitation pulse trigger and the photon arrival time (Datta et al., 2020) (Figure 3A). The result of the measurement is the distribution of photon arrival times. This is typically an exponential decay, which must be analyzed to extract a lifetime estimator (Figure 3B).

**Figure 3:**
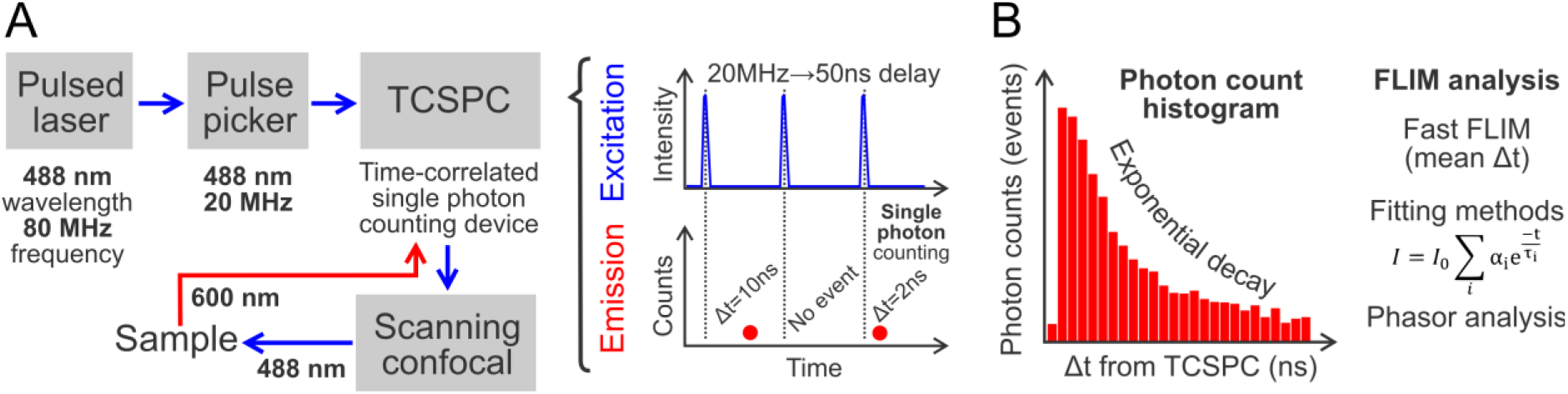
Principle of Fluorescence Lifetime Imaging Microscopy (FLIM). (**A**) Example of Flipper-TR FLIM set-up. A blue pulsed laser (488 nm, 80 MHz) is complemented with a pulse picker to reduce the frequency to 20 MHz (i.e., one pulse every 50 ns) allowing to capture the full decay. A time-correlated single photon counting device (TCSPC) times the delay between each pulse (blue) and each photon counting event (red dots). A scanning confocal allows for sample acquisition and optimize acquisition settings. (**B**) The output of the FLIM is histogram of photon arrival times (red), showing an exponential decay. The distribution of photon arrival times should be analyzed using different methods: fast FLIM, fitting methods, or phasor analysis.

Generally, the fluorescence lifetime is on the order of nanoseconds (Lakowicz, 2007). The resulting time-dependent emission *I*(*t*) is often described using the exponential model: *I*(*t*) = ∑_*i*_ α_*i*_*e*^−*t*∕τ*i*^. This distribution of the photon arrival times at the detector can be analyzed to determine the fluorescence lifetime, represented as τ, and the contribution of each exponent represented by the coefficient α such as ∑_*i*_ α_*i*_ = 1. Due to the photochemistry of the molecule, Flipper-TR photon arrival time distribution follows a bi-exponential decay in the form: *I*(*t*) = *A*_1_*e*^−*t*Ττ1^ + *A*_2_*e*^−*t*Ττ2^, where *A*_1_, *A*_2_ and τ_1_, τ_2_ represent the amplitude and the decay of a given exponent, respectively. Previous studies have employed different methods to estimate the Flipper-TR lifetime. One common approach is to fit the fluorescence decay curve to a biexponential decay model, which accounts for the presence of two distinct exponential decay components (Colom et al., 2018). From this model, researchers can extract either the main or larger exponent (i.e.: τ_1_) or calculate an intensity-weighted average of the two exponents, τ_av int_ = ∑_*i*_ τ_*i*_α_*i*_ (Roffay et al., 2021). Fitting methods can provide detailed insights into the Flipper-TR lifetime, including a χ^2^ used to judge the goodness, a greater precision, and the ability to decompose complex lifetime distributions, but they can be more sensitive to noise and initial parameter estimates. Moreover, even at high photon counts, the ability to determine the precise values of α_*i*_ and τ_*i*_ by a biexponential fit can be hindered by parameter correlation (Lakowicz, 2007). Parameter correlation means that the fitting parameters (α_*i*_ and τ_*i*_) are not independent, so that changes in one parameter can be compensated by changes in another while producing a very similar fit. For example, in a biexponential model, slightly increasing α_1_ while decreasing τ_2_and adjusting α_2_and τ_2_may yield the same decay curve. This makes it difficult for the fitting algorithm to uniquely determine each parameter, especially in limited photon count conditions. That’s why in practice, although biexponential fits capture the Flipper-TR photophysics better, researchers often report the fits using a single intensity-weighted average lifetime τ_av int_. Fitting methods used to have a large disadvantage of being computationally costly, but this has become a minor factor with modern PCs.

To avoid problems involved with fitting, other works have instead relied on the barycenter of the lifetime distribution (i.e., mean photon arrival time or “fast lifetime” calculation) to estimate fluorescence lifetime (García-Arcos et al., 2024). As the instrumentation induces a delay in the photon arrival, the average lifetime equals time span from the barycenter of the instrument response function (IRF) to the barycenter of the decay in a pixel-wise manner, although this definition depends on the TCSPC (Time-Correlated Single Photon Counting) manufacturer (Leica systems time both the pulse and the photon arrival times, so the IRF calculation is not needed for fast lifetime). When averaging or binning pixels, the lifetimes of each pixel τ_*i*_ are then weighted by the photon count *n*_*i*_, so that τ_fast_ = [∑^*n*^_i=1_ (*n*_*i*_τ_*i*_)]Τ*N*. This method is simple, straightforward, and computationally efficient. The calculation yields a single average Flipper-TR lifetime value, which we refer to as “fast lifetime” or τ_fast_, numerically equivalent to the intensity-weighted average of the two exponents coming from a fit. The need for such robust lifetime estimator at low photon counts is important when measurements are limited by sample illumination.

Finally, phasor plots are another fast and fit-free method recently used to estimate changes in Flipper-TR lifetime (Puji Pamungkas et al., 2024; Ragaller et al., 2024b). Phasor plots offer a graphical representation of the fluorescence decay characteristics, simplifying the analysis of complex decay patterns. In this representation, each decay curve is mathematically transformed into a point with coordinates G (the real part) and S (the imaginary part) of the Fourier transform at the laser modulation or pulse frequency, which together define the phasor position. By mapping each pixel to specific G/S coordinates on the phasor plot, one can visually assess changes in membrane tension and identify different lifetime components without assuming any feature of the fluorescence decay, such as the number of exponents. This is particularly useful when using complex samples or Flipper variants where the photochemical properties have not been fully characterized (Puji Pamungkas et al., 2024).

Despite the widespread use of Flipper-TR, no consensus exists on the optimal lifetime extraction method, and current publications vary in the analytical approaches they report. Furthermore, while each estimator is theoretically equally valid under ideal conditions, their behavior under realistic conditions varies (Leray et al., 2011; Starling et al., 2023). This includes (low) photon budgets, varying biological complexity, and instrumentation noise.

In this work, we systematically compare multiple strategies to estimate Flipper-TR lifetime across diverse biological samples and three different instrumental setups (two scanning confocal microscopes from Leica and Nikon/Picoquant and a 2-photon FLIM from Leica). We examine how different analysis methods behave as a function of photon count and other acquisition parameters, and describe a theoretical framework to understand the statistical limits of lifetime precision in each case. Building on principles of photon counting and fluorescence decay statistics, we show that the standard deviation of lifetime estimates generally scales with the number of photons *N* as σ ∝ 1 ∕ √*N*, a universal behavior derived from the Fisher information (Köllner & Wolfrum, 1992). We further incorporate biological variability into this model, separating the cell-to-cell biological heterogeneity from the instrumental shot noise, which becomes negligible at high photon counts. This allows us to formulate practical guidelines for acquisition and analysis: how many photons are needed to detect a given biological difference in lipid packing, depending on the method used. By combining theoretical analysis, empirical benchmarking, and practical workflows, this work provides a comprehensive guide to FLIM analysis of Flipper dyes, enabling researchers to make informed decisions depending on the experimental context.

## 2. Results

### 2.1 Effect of estimator type and number of exponents on fluorescence lifetime

Throughout this study, we chose to work with two samples with comparable average lifetime but different characteristics: a standard 1mM fluorescein sodium salt solution on PBS buffer, displaying a monoexponential lifetime of 4.05 ns (Hammer et al., 2005), and C2C12 cells stained with Flipper-TR, displaying a bi-exponential lifetime ranging from 2-6 ns. In some cases, this was complemented with pure lipid preparations stained with Flipper-TR: i) GUVs (Giant Unilamellar Vesicles) made with a variety of lipids displaying different degrees of lipid order and ii) supported lipid bilayers (SLBs) as described in (García-Arcos et al., 2024). Pure lipid samples stained with Flipper-TR display a bi-exponential lifetime while having a spatially homogeneous distribution and low data dispersion similar to dye solutions.

To compare different FLIM analysis strategies, we applied to fluorescein samples (Figure 4) and C2C12 cells stained with Flipper-TR (Figure 5) the range of estimators available on the NIS/Symphotime software: fast lifetime (4A, 5A), n-exponential reconvolution (4B-C, 5B-C), and n-exponential tail fitting (4D-E, 5D-E). For each of these estimators, we extracted a pixel-wise calculation (left panels), a histogram of photon counts per lifetime, separating the intensity of each component in multi-exponential fits (middle panels), and semi-log plots of the decay histograms overlayed with the fits (right panels), showing the lifetime decay (Decay, green), the software-calculated instrument response function (IRF, blue), the fits (Fit, red), together with the fit residuals (purple, with shown chi-square statistic). Please note that, from a theoretical standpoint, fluorescein lifetimes are described by a mono-exponential model, whereas Flipper-TR lifetimes are characterized by a bi-exponential model due to the presence of two distinct decay components. However, we considered for both samples mono-exponential and bi-exponential fits to test the response of the estimators to non-ideal conditions, i.e., too many or too few exponents.

**Figure 4:**
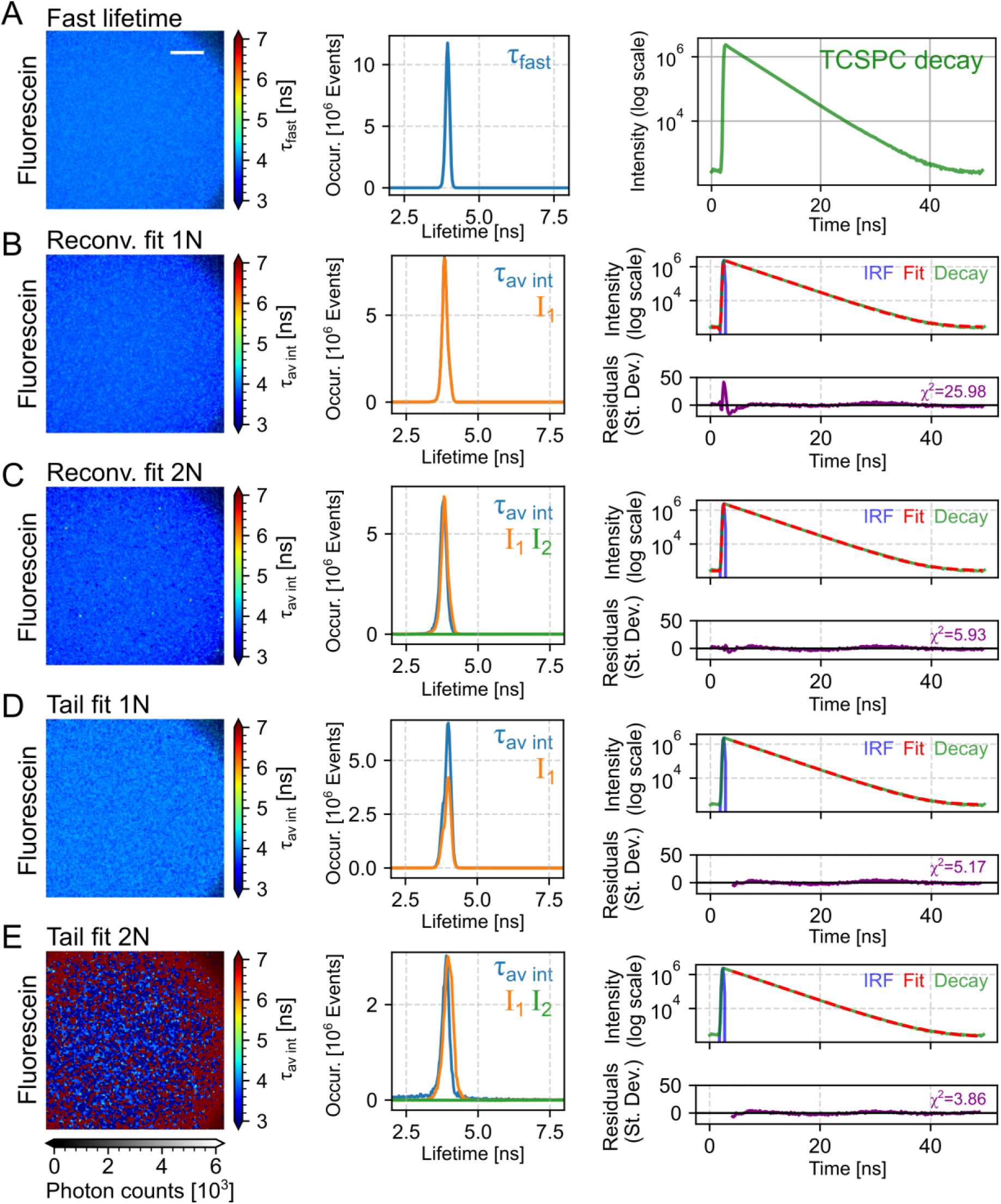
Fluorescence lifetime analysis of fluorescein standards. Data were acquired at a Nikon Ti2-A1R equipped with Picoquant FLIM hardware and analyzed using three different fitting approaches. (**A**) Fast lifetime map, showing pixel-wise lifetimes color-coded. A grayscale overlay, inverted and scaled by photon count, is used to visualize signal intensity. The central panel displays the distribution of lifetimes across pixels, and the right panel shows the corresponding time-correlated single photon counting (TCSPC) decay on a semi-logarithmic scale. (**B-E**) Results from pixel-wise fitting using single- or bi-exponential models, either by reconvolution (**B-C**) or tail fitting (**D-E**), as performed in SymPhoTime. Each panel includes: a lifetime map based on the fitted intensity-weighted average lifetime (left); a histogram of fitted intensity-weighted average lifetimes τ_av int_ and associated amplitude components I₁ and I₂ (center); and the TCSPC decay curve (green) overlaid with the instrument response function (IRF, blue) and fitted decay model (red dashed) (right). Residuals of the fit, expressed in standard deviations, are shown below the decay curves, and the χ^2^ value is indicated as a measure of goodness-of-fit. Scale bar, 50 µm.

**Figure 5:**
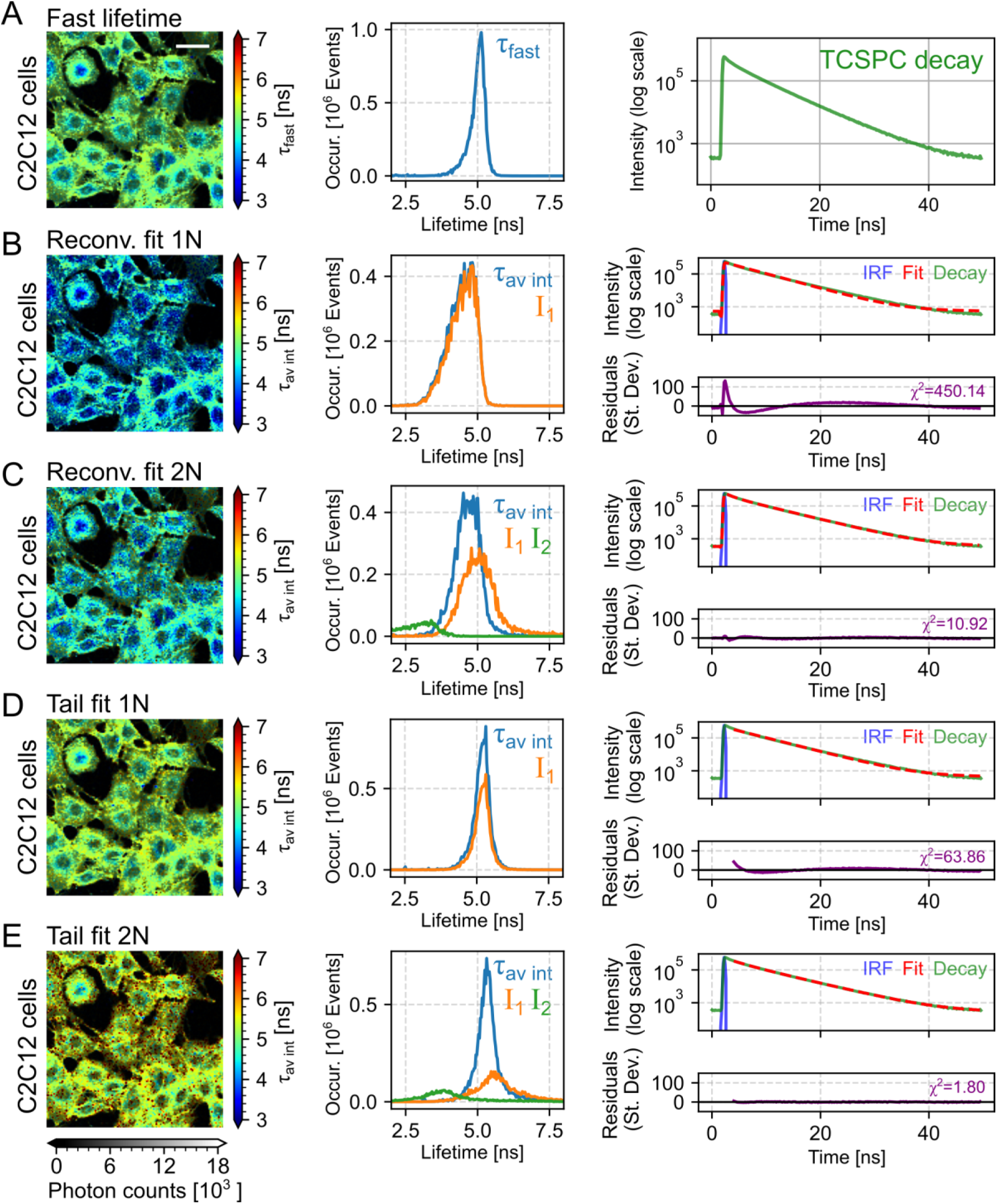
Fluorescence lifetime analysis of live C2C12 cells stained with Flipper-TR. Data were acquired at a Nikon Ti2-A1R equipped with Picoquant FLIM hardware and analyzed using three different fitting approaches. (**A**) Fast lifetime map, showing pixel-wise lifetimes color-coded. A grayscale overlay, inverted and scaled by photon count, is used to visualize signal intensity. The central panel displays the distribution of lifetimes across pixels, and the right panel shows the corresponding time-correlated single photon counting (TCSPC) decay on a semi-logarithmic scale. (**B-E**) Results from pixel-wise fitting using single- or bi-exponential models, either by reconvolution (**B-C**) or tail fitting (**D-E**), as performed in SymPhoTime. Each panel includes: a lifetime map based on the fitted intensity-weighted average lifetime (left); a histogram of fitted intensity-weighted average lifetimes τ_*av int*_ and associated amplitude components I₁ and I₂ (center); and the TCSPC decay curve (green) overlaid with the instrument response function (IRF, blue) and fitted decay model (red dashed) (right). Residuals of the fit, expressed in standard deviations, are shown below the decay curves, and the χ^2^ value is indicated as a measure of goodness-of-fit. Scale bar, 50 µm.

Fluorescein samples display a spatially homogeneous lifetime, as expected from a dye solution. The fast lifetime histogram is Gaussian-shaped and narrow (Figure 4A). The semi-log decay plot displays a stereotypical straight line, indicating that a single exponent is involved (4A, right panel, TCSPC decay). We then analyzed this decay using fitting algorithms, n-exponential reconvolution, and tail fitting. Reconvolution fits the entire measured fluorescence decay, convolved with the system’s instrument response function (IRF), to an exponential model, using the complete decay data. A proper measurement of the IRF might be important for this method if the computation estimation is not sufficient to achieve a good fit. Tail fit is a simpler approach considering only the latter portion (the “tail”) of the fluorescence decay curve. This can be a viable option when fluorescence lifetimes are significantly longer than the instrument response function. The 1-exponent reconvolution of fluorescein data shows a significant error arising from the segment surrounding the IRF, which results in a very short component appearing and a large chi-square (Figure 4B). This could be technically improved with an actual measurement of the IRF instead of relying on a computational estimation (e.g., with a fluorescein solution quenched on a saturated potassium iodide). Comparatively, the 1-exponent tail fit of fluorescein, relying less on the IRF estimation, achieves a better result and a lower fitting error (Figure 4D). When we attempted to fit fluorescein data with 2-exponential models, we observed that introducing additional exponential terms only increased the noise level without significantly improving the accuracy of the fit (as seen in the pixel-wise fit images on 4C, 4E). This is because the contribution of the additional lifetime component is extremely small and carries negligible weight in the overall lifetime histogram (4C, 4E green lines corresponding to I2). Consequently, multi-exponential fitting is unnecessary for fluorescein, and the mono-exponential model is the best approach.

Following the characterization of the fluorescein standard, we applied the same analytical framework to C2C12 cells stained with Flipper-TR. C2C12 cells display a spatially heterogeneous fast lifetime. The fast lifetime histogram is left-skewed and wider than the fluorescein case (Figure 5A). The semi-log decay plot displays a slightly curved line, indicating that more than one exponent is likely involved (5A, right panel, TCSPC decay). Both the 1-exponent reconvolution and tail fit of cells show a very significant error arising from the misestimation of the extra component, which is visible in the structure of the residuals (Figures 5B, 5D). The 2-exponent fits improve significantly the fitting errors for Flipper-TR-stained cells and is therefore the best approach (Figure 5C, 5E).

Increasing the number of decay exponents does not have the same impact on the fitting errors in the case of a single decay as fluorescein or in a more complex decay pattern, such as Flipper-TR on a biological sample. After confirming that a bi-exponential fitting of Flipper-TR fluorescence decay improves the overall goodness of fit, we next tested how the estimated lifetime values are affected by the number of exponents. We chose two indicators: τ_1_ (Tau 1, the longest lifetime component) and τ_av int_ (intensity-weighted average lifetime) under different exponential fitting models for both fluorescein and Flipper-TR, along with their corresponding errors. Note that the nomenclature for τ₁ can vary depending on the software used. In this work, τ₁ refers to the dominant lifetime component, the component with the largest weight.

For fluorescein standards, both τ_av int_ and τ_1_ tend to remain stable regardless of the number of exponents, except in cases where increasing the number of exponents introduces additional error (Figure 6A). This effect is consistent with the single-field analysis shown in Figure 4. The three-exponential fit produced poorer results than the bi-exponential fit, particularly using tail fitting (Figure 6B). In cells stained with Flipper-TR, we observed that adding more exponential components consistently increased both τ_av int_ and τ_1_, and reduced the fitting error. Cells display a complex decay pattern, and additional shorter components take a significant portion of the overall decay amplitude, causing τ_1_ to increase its value to compensate. As a consequence, the values of τ_av int_, the average between all components, fluctuate less with increasing number of exponents (Figure 6C-D). This indicates that in cells, at 3 exponents, the model is not yet overparametrized (i.e., many more parameters or degrees of freedom than needed). This is probably due to the presence of residual lifetime components within the cytoplasm, as well as possible contributions from non-specific staining or cellular debris. Unlike fluorescein, where a single-exponential fit is sufficient, for cells, at least 2 exponents must be used.

**Figure 6:**
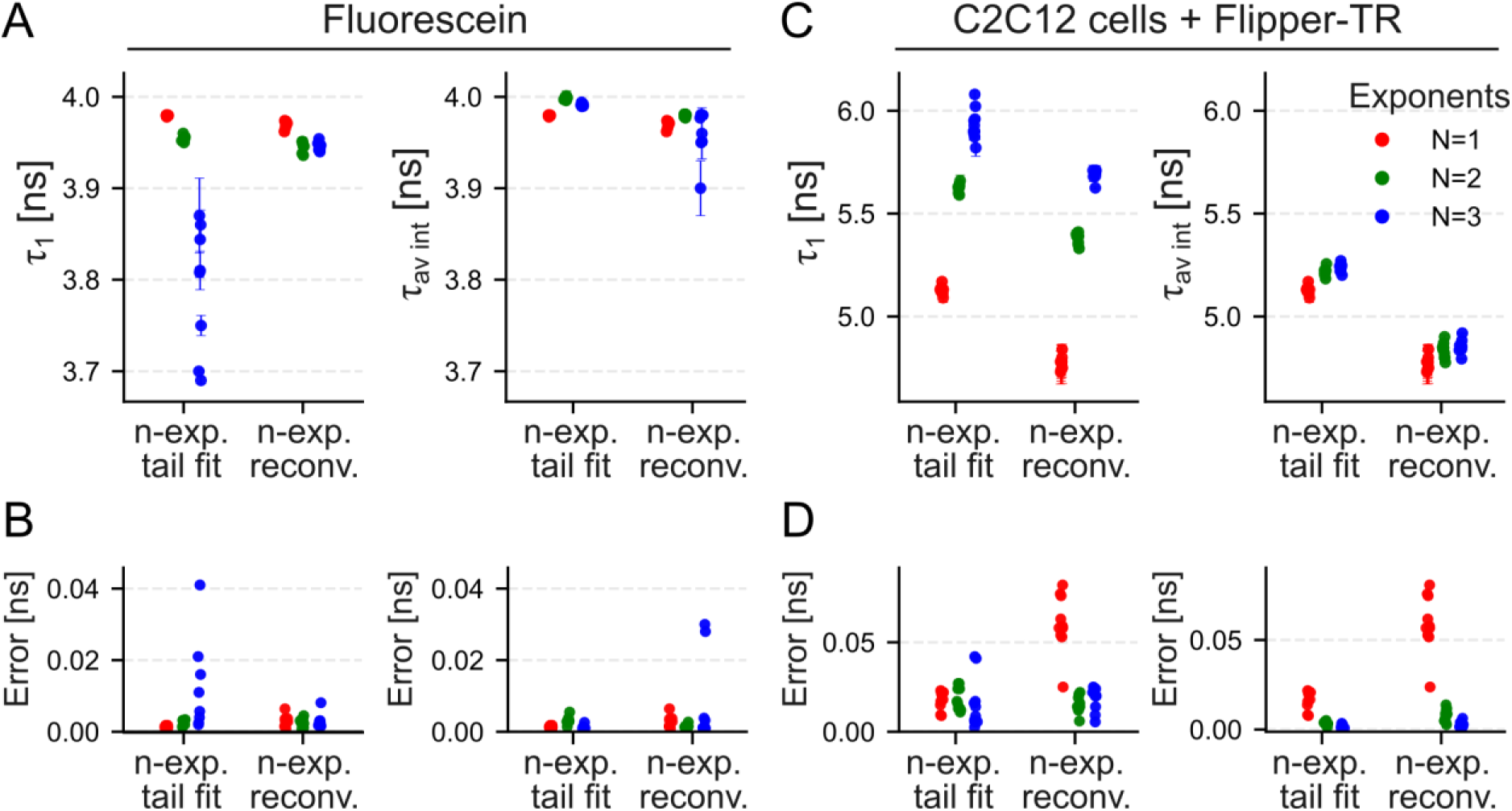
Component lifetimes and reported fit uncertainty by fitting strategy for C2C12 cells and fluorescein standards. (**A–B**) Short component lifetime (τ_1_) and intensity-weighted average lifetime (τ_*av int*_) grouped by fit method (tail fit, reconvolution). Columns show τ_1_ (left) and τ_*av int*_ (right) for C2C12 (A) and Fluorescein (B). Points are individual measurements; vertical bars are ±Err reported by SymPhoTime. Colors encode the number of exponential components used (N: red = 1, green = 2, blue = 3). Axes are in nanoseconds. (**C-D**) Corresponding distributions of ±Err shown alone for the panels above (A-B), with identical x-grouping and color coding.

Overall, τ_av int_ values tend to be more stable than τ_1_. This is because τ_av int_ integrates information across all exponentials, reflecting the overall decay shape, which is a robust property of the decay curve itself.. On the other hand, τ_1_ can take more extreme values as long as other lifetime components compensate for this. This is especially true in cases where the model is over-parametrized. To further evaluate which fitted lifetime parameters provide consistent and reliable readouts, we compared the correlation between τ_av int_, τ_1_ and τ_2_ from 2-exponential reconvolution fits with ideal imaging conditions (photon counts per fit > 10^5^; Signal-to-noise ratio > 20). This analysis allows us to assess whether the main component and the intensity-weighted average lifetime capture similar environmental sensitivity, and how they relate to the secondary component in terms of reproducibility and fitting uncertainty. We focused on pure lipid samples (GUVs) of 6 different compositions stained with Flipper-TR. This provided a range of lifetime values while eliminating the spatial heterogeneity present in cell samples. Each fit in the plot corresponds to an individual field of view containing several GUVs (Figure 7). In cases where the composition was phase-separating into liquid-ordered (Lo) and liquid-disordered (Ld) domains, we segmented and fitted them independently.

**Figure 7:**
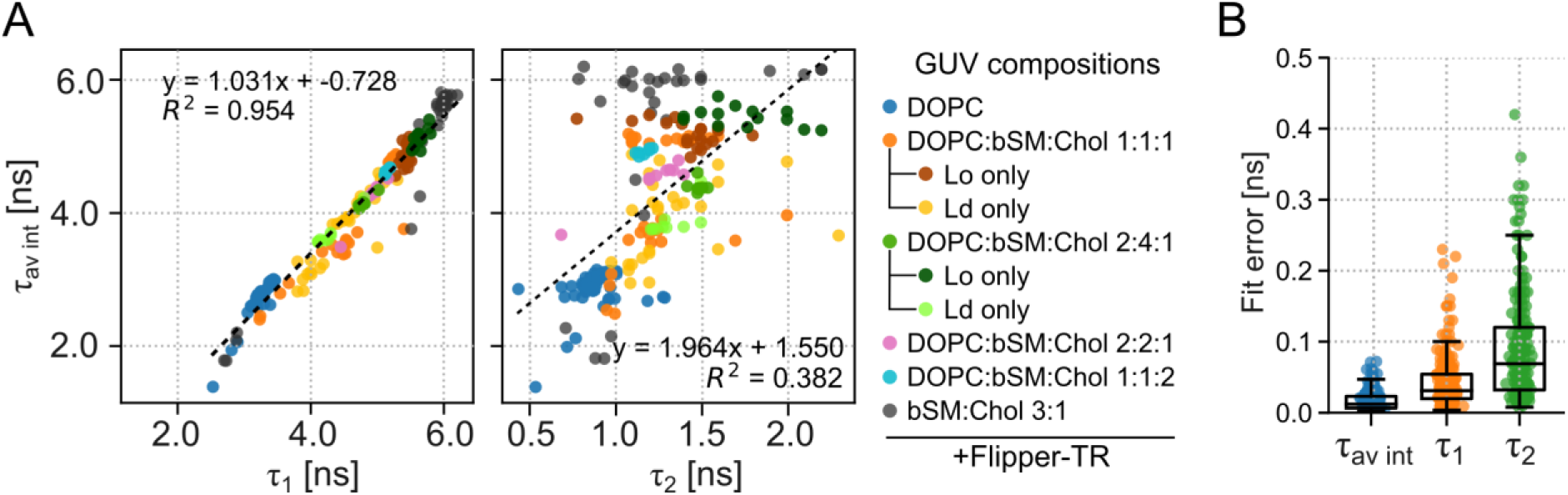
Correlation of component lifetimes with intensity-weighted average lifetime across GUV compositions. (**A**) Left, scatter plot of the longest lifetime component (τ_1_) versus the intensity-weighted average lifetime (τ_av int_) for individual GUV measurements using n-exponential reconvolution fitting on Symphotime (Picoquant). Points are color-coded by lipid composition: DOPC; DOPC:brain sphingomyelin:cholesterol (1:1:1), (4:2:1), and (2:2:1); and brain sphingomyelin:cholesterol (3:1). For compositions exhibiting phase separation, lifetimes measured within regions assigned to liquid-ordered (Lo) or liquid-disordered (Ld) phases and plotted as separate data points. Right, minor lifetime component (τ_2_) versus τ_av int_. A least-squares linear fit is overlaid; the regression equation and coefficient of determination (R²) are displayed. (**C**) Distribution of fit uncertainties (±Err) reported by the analysis on Symphotime for τ_1_, τ_2_, and τ_av int_ across all conditions. Box-and-whisker plots represent median and interquartile range (boxes), with whiskers to 1.5×IQR; individual values are overlaid as points.

Our analysis revealed that τ_av int_ values show a clear linear correlation with τ₁, with a slope ≈1 and a shift of ≈0.73 nanoseconds. In contrast, τ_2_ values did not exhibit any correlation with τ_av int_ intensity (Figure 7A). This is consistent with the mechanism proposed in other works: τ₁ is linked to the time taken for the emission from the PICT state, therefore linked to the mechanism by which Flipper-TR senses lipid packing. From these observations, we conclude that τ₂ is not a reliable estimator of Flipper-TR lifetimes. Instead, only τ_av int_ and τ₁ are indicators of Flipper-TR conformation.

Due to the biexponential fitting algorithm, τ_av int_ displays less dispersion than τ_1_. In a bi-exponential fit, τ_1_, τ_2_ and their amplitudes are strongly anti-correlated: when τ_2_jitters up, its fitted amplitude tends to jitter down, and τ_2_shifts the opposite way (Lakowicz, 2007). This can be intuitively understood from the variance propagation of two anti-correlated variables. If τ_av int_ = *A*_1_τ_1_ + *A*_2_τ_2_; and we locally treat *A*_1_ and *A*_2_ as fixed weights, then Var(*t*_av int_) ≈ *A*^2^Var(τ_1_) + *A*^2^Var(τ_2_) + 2*A*_1_*A*_2_Cov(τ_1_, τ_2_). Because *A*_2_ is small and τ_1_ ∝ −τ_2_, then the term Cov(τ_1_, τ_2_) < 0 and subtracts variance. For this reason, the fit errors appear smaller in τ_av int_ compared to individual components (Figure 7B). From this, we conclude that τ_av int_ a more robust estimator for Flipper-TR lifetime than any of the individual components.

### 2.2 Effect of microscopy setup and laser frequency on Flipper-TR lifetime estimators

So far, we have explored the influence of data analysis from ideal datasets from a single microscope setup. Practically, users might be constrained to acquire data in non-ideal conditions or use a variety of setups with different characteristics. We next explored the effect of the microscopy setup and laser frequency on the different Flipper-TR estimators. For this, we compared the average lifetime of Flipper-TR-stained C2C12 cells and the fluorescein standard in ideal conditions using three different scanning confocal FLIM systems: a Nikon-Picoquant integration with Sepia PDF 828 laser and MultiHarp 150 card with PMA Hybrid 40 detector using Symphotime 64 software for analysis, a Leica Stellaris Falcon with HyD X detector, and a Leica 2-photon SP8 DIVE with external super HyD detector, both using LAS X software for analysis (Table 1). Both systems reply of TCSPC for estimating the lifetime, but have their own proprietary algorithms and some differences in the hardware. In Leica LAS X, both the excitation pulse and the photon arrival time are recorded, allowing the software to directly calculate the delta-time for each photon relative to the laser pulse. This approach means that the instrument response function (IRF) is inherently considered, so the fitted IRF shift is typically close to zero. In contrast, in Nikon NIS systems using PicoQuant hardware, only the photon arrival times are recorded, and the IRF must be separately estimated. In this case, the fast lifetime component is obtained from the difference between the barycentre of the IRF and the decay curve, which introduces an additional source of uncertainty and makes the fitted IRF shift more prominent.

**Table 1:** average fluorescein standard lifetime and C2C12 cells stained with Flipper-TR in different experimental set-ups. Data is reported as mean ± standard deviation between samples for at least N>3.

| | Microscope<br>(scanning confocal) | Pulse<br>frequency<br>(MHz) | Fast lifetime<br>$\tau_{\text{fast}}$ (ns) | Intensity-weighted average<br>lifetime $\tau_{\text{av int}}$ (ns) | |
| --- | --- | --- | --- | --- | --- |
|  |  |  |  | n-exponent<br>reconvolution fit | n-exponent<br>tail fit |
| Fluorescein | *Nikon-Picoquant Ti2 A1* | 20 | 3.91 ± 0.06 | 3.97 ± 0.00 | 3.98 ± 0.00 |
|  | *Leica Stellaris Falcon* | 20 | 3.97 ± 0.05 | 3.99 ± 0.00 | 3.99 ± 0.00 |
|  | *Leica 2-photon SP8 DIVE* | 80 | 3.26 ± 0.00 | 4.13 ± 0.02 | 4.14 ± 0.02 |
| Flipper-TR | *Nikon-Picoquant Ti2 A1* | 20 | 4.90 ± 0.06 | 4.84 ± 0.04 | 5.22 ± 0.02 |
|  | *Leica Stellaris Falcon* | 20 | 5.30 ± 0.10 | 5.12 ± 0.03 | 5.20 ± 0.02 |
|  | *Leica 2-photon SP8 DIVE* | 80 | 3.34 ± 0.02 | 4.40 ± 0.17 | 4.52 ± 0.11 |

Surprisingly, the lifetime values of Flipper-TR exhibit significant variability across different microscopes. This suggests that comparing or pooling absolute lifetime values acquired from different microscope set-ups is not advisable, in particular across instruments from different manufacturers. Note that the trends between the different estimators τ_fast_, τ_av int_ from reconvolution and τ_av int_ from tail fits also differ between setups, suggesting that the behavior of these estimators might not be the same in different setups despite bearing the same name. Generally, the lifetime values that differ the most are those obtained from our commercial two-photon microscope, which surprisignly also appear for the fluorescein standard. This is likely due to the repetition rate, which is set at 80 MHz for our 2-photon laser. In TCSPC-based FLIM, for the determination of the absolute lifetime values, the interval between excitation pulses must cover most of the decay of the fluorophore to avoid temporal overlap between successive decay curves. However, at 80 MHz, the pulse interval is only 12.5 ns, which is insufficient for complete relaxation of fluorophores with longer lifetimes such as our fluorescein standard and the Flipper-TR-stained cells. This results in a distortion of the fluorescence decay profile, particularly an underrepresentation of the long-lifetime components, ultimately leading to inaccurate lifetime estimation. Conversely, the 20 MHz laser, with a 50 ns pulse interval, provides sufficient time for full fluorescence decay before the arrival of the next pulse, enabling correct reconstruction of the full decay. These constraints cannot be avoided in some setups, as most commercial 2-photon lasers cannot modulate the repetition rate, set at 80 MHz.

To explore the effect of laser repetition rate on lifetime estimation, we compared a fluorescein standard (Figure 8) and C2C12 cells stained with Flipper-TR (Figure 9) acquired using pulsed laser sources at 80, 40, and 20 MHz on our Nikon/Picoquant setup. For each of these, we extracted: i) a pixel-wise fast lifetime τ_fast_ and a histogram of τ_fast_ values, ii) a n-exponential reconvolution fit, IRF, and residuals, and iii) the equivalent for the tail fits. The number of exponents for fluorescein and Flipper-TR cells were set at 1 and 2, respectively. For fluorescein, the fast lifetime values highly depend on the laser repetition rate (see false-color images on Figure 8A). Because the fast lifetime only records the photon arrival time, shortening the window of acquisition has a strong effect on the center-of-mass of the fast lifetime histogram (see histogram on Figure 8A). The fits revealed that the 1-exponential reconvolution τ_av int_ is highly sensitive to variations in the laser repetition rate, and fits cannot converge well with the data at low repetition rate (Figure 8B, top row). The fit does not take into account the effect of the shortened decay window before the IRF. The fit is flat before the IRF, while the data shows a mini decay. In contrast, the tail-fitting method demonstrated great robustness, exhibiting no sensitivity of τ_av int_ to changes in the laser repetition rate, and low fit residuals (Figure 8B, bottom row). Note that this constraint is imposed by the specific fitting algorithm and may vary in other software tool and manufacturer.

**Figure 8:**
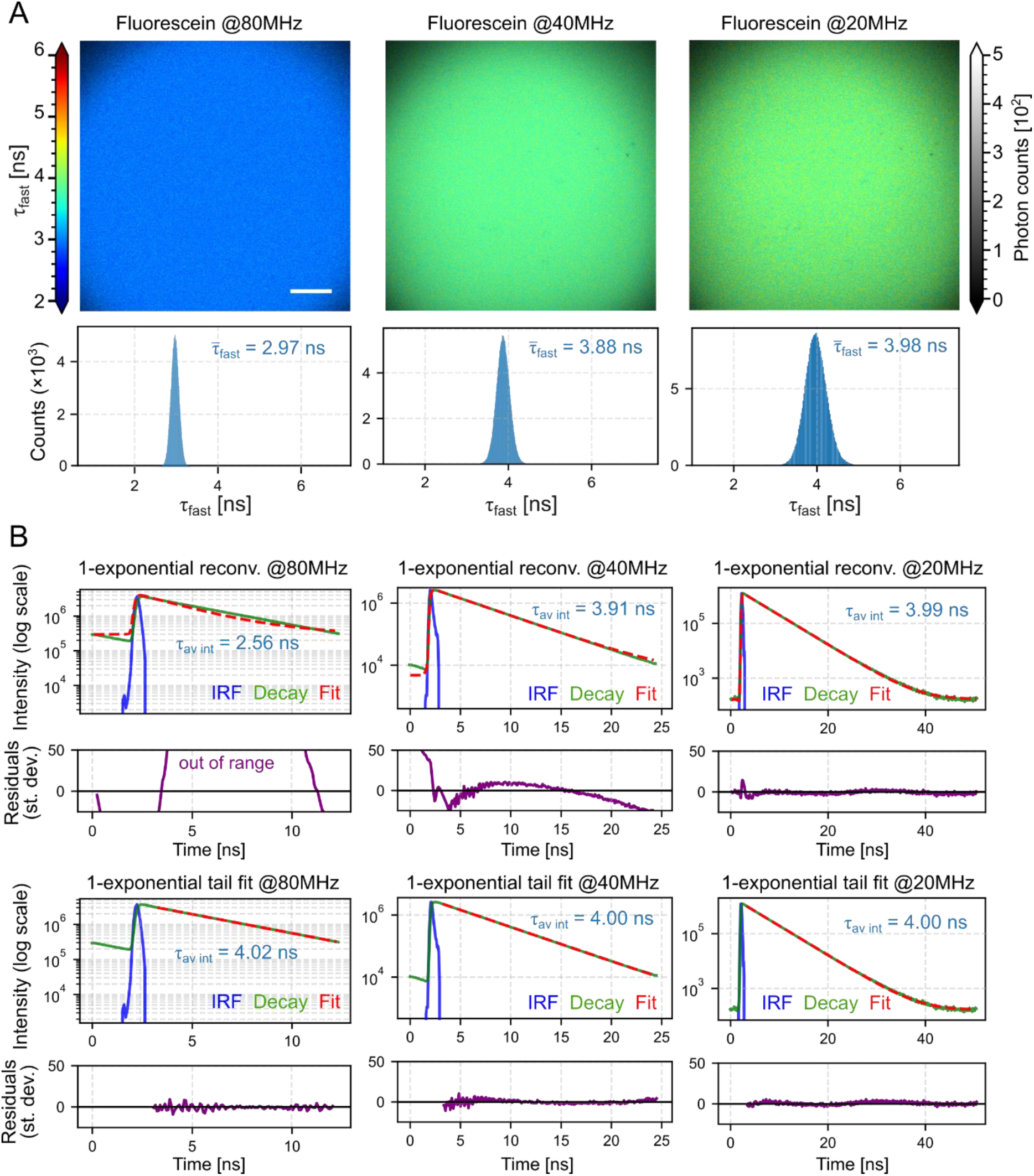
Frequency dependence of fast-lifetime FLIM and fits for fluorescein. (**A**) Fast lifetime images and distributions at three laser repetition rates. Columns correspond to excitation frequencies 80 MHz (left), 40 MHz (middle), and 20 MHz (right). Top row: fast-lifetime maps (τ_fast_) with grayscale intensity overlay; the color bar reports τ_*fast*_ in ns. Bottom row: histograms of τ_fast_ for the same fields of view; the mean fast lifetime (τ_fast_) is shown as a blue annotation on each histogram. (**B**) TCSPC decays and fitting residuals for the dame dataset as (A). First two rows: 1-exponential reconvolution (fit shown as red dashed). Traces are IRF (blue), decay (green), and fit (red dashed). Blue label reports τ_av int_. The row below each fit panel shows the residuals (purple) in units of standard deviations versus time (ns). Last two rows: same as in top row for tail fitting. Scale bar, 50 µm.

**Figure 9:**
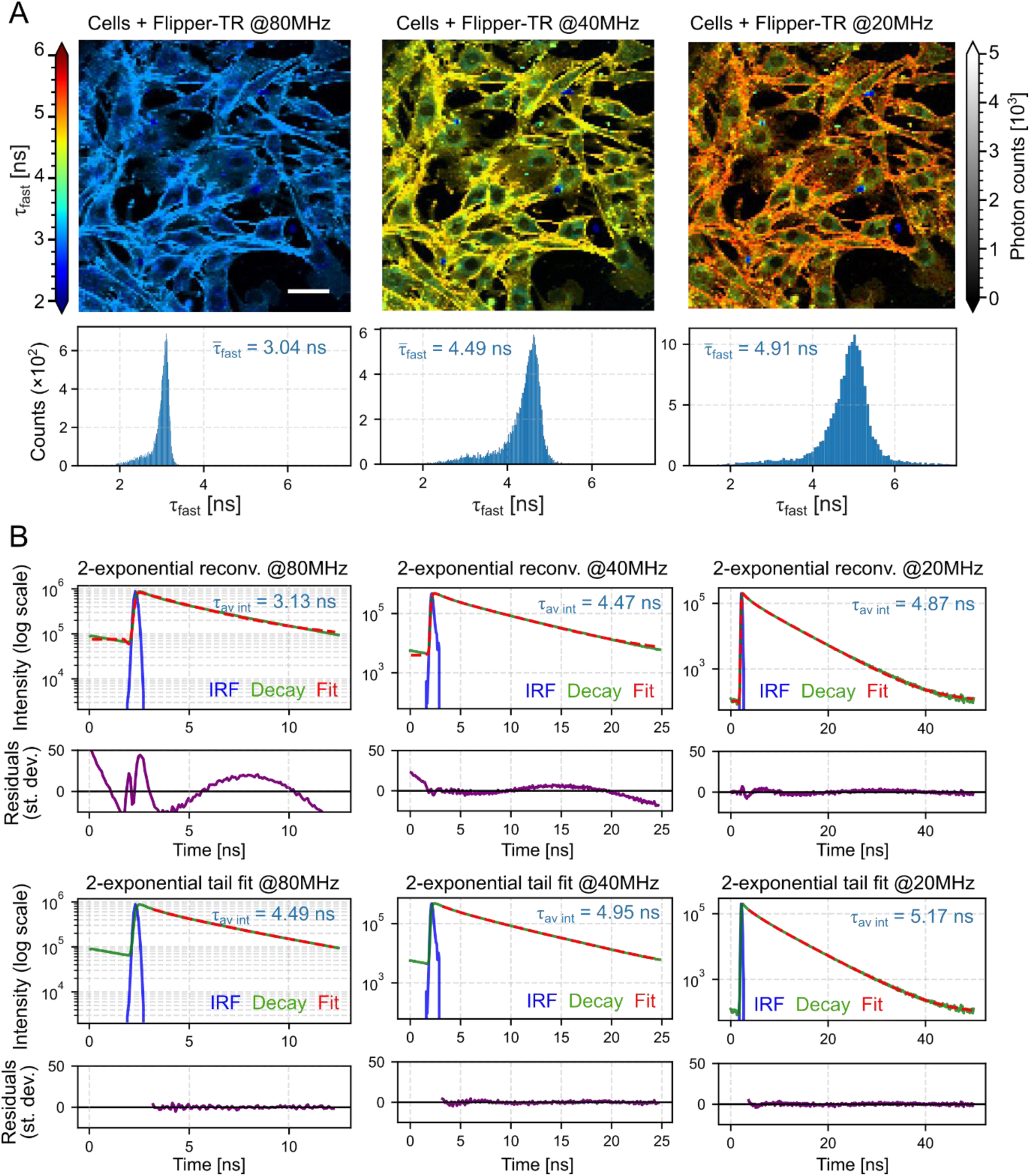
Frequency dependence of fast-lifetime FLIM and fits for C2C12 cells. (**A**) Fast lifetime images and distributions at three laser repetition rates. Columns correspond to excitation frequencies 80 MHz (left), 40 MHz (middle), and 20 MHz (right). Top row: fast-lifetime maps (τ_fast_) with grayscale intensity overlay; the color bar reports τ_*fast*_ in ns. Bottom row: histograms of τ_fast_ for the same fields of view; the mean fast lifetime (τ_fast_) is shown as a blue annotation on each histogram. (**B**) TCSPC decays and fitting residuals for the dame dataset as (A). First two rows: 2-exponential reconvolution (fit shown as red dashed). Traces are IRF (blue), decay (green), and fit (red dashed). Blue label reports τ_av int_. The row below each fit panel shows the residuals (purple) in units of standard deviations versus time (ns). Last two rows: same as in top row for tail fitting. Scale bar, 50 µm.

The results for Flipper-TR-stained cells agree qualitatively, indicating that both the fast lifetime τ_fast_ (Figure 9A) and the bi-exponential reconvolution τ_av int_ estimators are highly sensitive to changes in the laser repetition rate (Figure 9B, top row). In contrast, the bi-exponential tail τ_av int_ showed lower sensitivity to variations in laser frequency (Figure 9B, bottom row). However, unlike with the fluorescein standard solution, the bi-exponential tail fit τ_av int_ still underestimates the absolute lifetime value of Flipper-TR at 40 and 80 MHz laser frequencies. Therefore, in cells stained with Flipper-TR, no estimator is completely insensitive to changes in the laser pulse frequency.

The measurement of the instrument response function by a quenched dye (in this case 200mM fluorescein sodium salt on a saturated solution of potassium iodide) can help with the accuracy of the n-exponential reconvolution fits at a high frequency (Table 2), but the tail fits as expected are not affected by changes in the IRF. In ideal conditions, at 20MHz, our quenched dye solution actually performs worse than the software-generated one in performing fits.

**Table 2:** Intensity-weighted average lifetimes, errors and χ^2^ from fits at different frequencies (20,40, 80 MHz) and using a measured IRF (quenched fluorescein on KI) or a software-generated IRF.

| Sample | Frequency | Estimator | IRF type | $\tau_{\text{av int}}$ [ns] | Error (±) | $\chi^2$ |
| --- | --- | --- | --- | --- | --- | --- |
| Fluorescein standard | 20MHz | Reconv. | Measured | 3.96 | 0 | 44.943 |
|  |  |  | Software | 3.99 | 0 | 2.395 |
|  | 40MHz |  | Measured | 3.951 | 0 | 31.948 |
|  |  |  | Software | 3.902 | 0 | 50.201 |
|  | 80MHz |  | Measured | 3.9 | 0 | 63.195 |
|  |  |  | Software | 3.25 | 0 | 719.662 |
|  | 20MHz | Tail fit | Measured | 4.001 | 0 | 2.197 |
|  |  |  | Software | 4.001 | 0 | 2.197 |
|  | 40MHz |  | Measured | 3.982 | 0 | 1.356 |
|  |  |  | Software | 3.982 | 0 | 1.356 |
|  | 80MHz |  | Measured | 3.972 | 0 | 3.094 |
|  |  |  | Software | 3.97 | 0 | 3.094 |
| C2C12 cells + Flipper-TR | 20MHz | Reconv. | Measured | 4.84 | 0.009 | 53.239 |
|  |  |  | Software | 4.88 | 0.01 | 24.184 |
|  | 40MHz |  | Measured | 4.6 | 0.01 | 227.003 |
|  |  |  | Software | 4.54 | 0.005 | 320.649 |
|  | 80MHz |  | Measured | 4.03 | 0.014 | 106.405 |
|  |  |  | Software | 3.12 | 0.01 | 1158.279 |
|  | 20MHz | Tail fit | Measured | 5.354 | 0.026 | 1.284 |
|  |  |  | Software | 5.354 | 0.023 | 1.284 |
|  | 40MHz |  | Measured | 5.08 | 0.001 | 3.68 |
|  |  |  | Software | 5.077 | 0.001 | 3.68 |
|  | 80MHz |  | Measured | 3.972 | 0 | 3.094 |
|  |  |  | Software | 3.97 | 0 | 3.094 |

**Table 2:** Summary of the analysis strategies benchmarked in this study. Columns indicate whether fitting is required, dependence on the instrument response function (IRF), sensitivity to laser repetition rate, precision under photon-limited conditions (low N), suitability for absolute quantification, and typical recommended use. †Definition is vendor-dependent; Leica timing obviates explicit IRF handling for fast lifetime.

| Flipper-TR lifetime estimator | $\tau$ fast | $\tau$ average (reconvolution) | $\tau$ average (tail fit) | $\tau_1$ (major component) | Phasor (G,S) |
| --- | --- | --- | --- | --- | --- |
| Fit? | No | Yes | Yes | Yes | No |
| IRF needed | No† | Yes (measured preferred) | Reduced need | As per fit | No |
| Behavior vs. laser rep. rate | High sensitivity | High sensitivity | Lower sensitivity than reconvolution | As per fit | — (model-free; plotted at $\omega$) |
| Precision at low photon counts N | High (best on Nikon; ≈ others on Leica) | Medium on Nikon; ≈ others on Leica | Lowest on Nikon; ≈ others on Leica | More variable than $\tau$ average | Robust directional shifts at very low N. Low variability. |
| Absolute quantification | Setup-dependent; avoid cross-instrument pooling | Main estimator when IRF reliable; prefer 20 MHz | Robust when IRF unreliable; not for freq. conversion | Mechanistic, not sole metric | Calibration needed for absolute. Hard to compare with any other estimator. |
| Typical use | Fast check, dynamics, photon-limited | Absolute lifetimes (with good IRF) | Alternative/validation | Supportive interpretation | Low-photon or complex sample, complementary |

To test how lifetime estimators behave across different laser repetition rates, we took advantage of the natural spatial heterogeneity of Flipper-TR in C2C12 cells (Figure 9A). Within the same cell, some regions show low lifetimes and others high lifetimes, which provides an internal reference range. We sequentially acquired the same fields of view at 20, 40, and 80 MHz. Because cell movement was negligible during these rapid acquisitions, this approach enabled a pixel-wise comparison of lifetime values across frequencies. To minimize noise, we acquired high photon counts (>10^5^; SNR>50) and used large pixels (5-10 µm). Our goal was to determine whether lifetime values at different repetition rates could be related by a simple linear conversion.

We compared three estimators on the Nikon/PicoQuant setup: the mean-arrival-time τ_fast_, the intensity-weighted average lifetime τ_av int_ from bi-exponential reconvolution, and τ_av int_ from bi-exponential tail fits. First, we compared each estimator across repetition rates at 20, 40, and 80 MHz (Figure 10). For each pair of estimator, we performed pixel-wise scatter plots and fitted a robust linear model (weighted Huber), reporting slope, intercept, and R^2^ on each panel. τ_fast_ (Figure 10A) and τ_av int_ from bi-exponential reconvolution (Figure 10B) exhibited clear linear relationships between frequencies, with R^2^ highest for 40 vs. 80 MHz, intermediate for 20 vs. 40 MHz, and lowest, although significant for 20 vs. 80 MHz (Figure 10A-B). By contrast, τ_av int_ from tail fit showed weak or absent linear correspondence across frequencies (Figure 10C).

**Figure 10:**
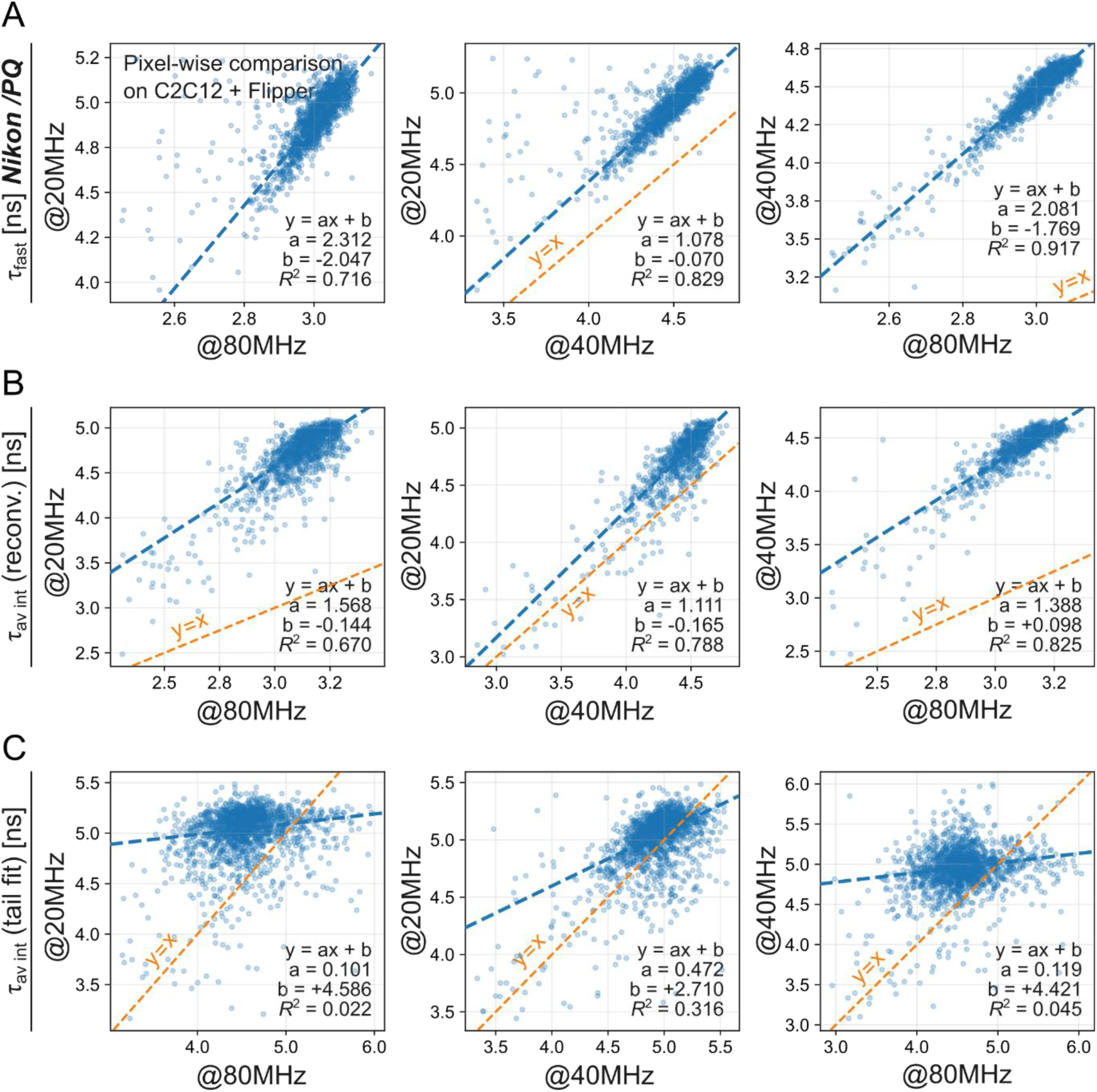
Correspondence between laser frequencies on Nikon/Picoquant. Rows (top to bottom): lifetime estimators τ_fast_, τ_av int_ (reconvolution), τ_av int_ (tail fit). Columns (left to right): pixel-wise scatter of 20 versus 80MHz; 20 versus 40MHz; and 40 versus 80MHz. Each dot is one pixel. Axes report lifetime (ns) with different estimators depending on the row. Points are overlaid with a weighted Huber regression, a robust linear fit (blue dashed) and a y=x identity line (orange dashed). Each panel annotates slope, intercept, and R². Inclusion filters: lifetime ∈ [2, 10] ns in both images; intensity > 1000 counts in both images; symmetric percentile clipping on lifetimes (1st–99th) applied after the hard thresholds.

We then re-organized the analysis to instead compare estimators against each other at fixed frequencies (Figure 11). τ_fast_ and τ_av int_ from bi-exponential reconvolution were linearly related at all three frequencies (Figure 11, first column). In contrast, τ_av int_ from tail fit was linearly related to τ_fast_ and τ_av int_ from reconvolution at 20 and 40 MHz (Figure 11A) and degraded at 80 MHz (Figure 11B-C). Together, these observations are consistent with τ_av int_ from tail fit being less sensitive to repetition-rate changes than the other two estimators, but also less suitable for establishing a frequency-to-frequency conversion on this setup.

**Figure 11:**
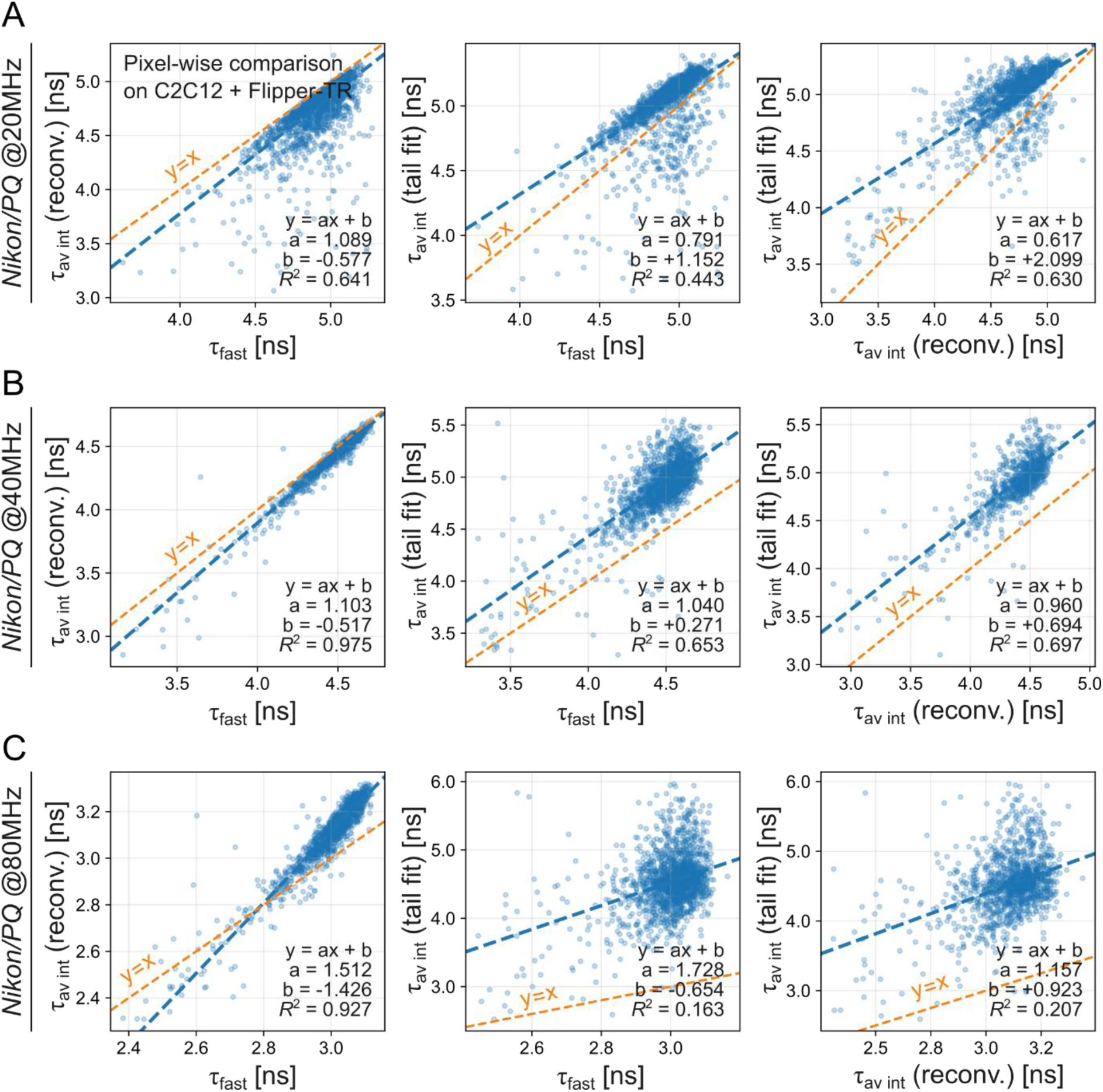
Correspondence between fit models on Nikon/Picoquant. Rows (top to bottom): laser repetition rates 20, 40, 80 MHz. Columns (left to right): pixel-wise scatter of τ_fast_ versus τ_av int_ (reconvolution); τ_fast_ versus τ_av int_ (tail fit), and τ_av int_ (reconvolution) versus τ_av int_ (tail fit). Each dot is one pixel. Axes report lifetime (ns) with different estimators as indicated on the axes. Points are overlaid with a weighted Huber regression, a robust linear fit (blue dashed) and a y=x identity line (orange dashed). Each panel annotates slope, intercept, and R². Inclusion filters: lifetime ∈ [2, 10] ns in both images; intensity > 1000 counts in both images; symmetric percentile clipping on lifetimes (1st–99th) applied after the hard thresholds.

Next, we aimed to investigate whether the conclusions drawn from our analysis comparing estimators and frequencies at Nikon/Picoquant would hold when using a different microscope setup. In Table 1, we present results showing that, even when analyzing the same sample at the same excitation frequency, the absolute lifetime values differ across microscope setups. This discrepancy arises because, although different manufacturers may label parameters similarly, the underlying hardware and data processing algorithms are not identical. To test this, we repeated the frequency-conversion analysis on a Leica system (Figures 12–13). Within the Leica dataset, pixel-wise comparisons analogous to Figure 10, organized as cross-frequency regressions (Figure 12), none of the estimators could provide a reasonable conversion between frequencies. When organized to test the relation between different estimators, the intensity-weighted average lifetime τ_av int_ from bi-exponential reconvolution, and τ_av int_ from bi-exponential tail fits were mutually correlated, whereas τ_fast_ did not track the fit-derived estimators consistently. At 20 MHz, all three estimators correlated reasonably well; however, correlations degraded at 40 and 80 MHz, particularly τ_fast_ versus the fit-derived estimators. Together, these results indicate that frequency-to-frequency calibration is setup-dependent: on this Leica setup, fitting lifetime estimates are less affected by the wrap-arounds on shorter pulses. This robust estimation disables a conversion between frequencies. Lifetime fitting is sensitive to specific processing to each software, therefore absolute values should not be pooled or converted across systems without instrument-specific calibration.

**Figure 12:**
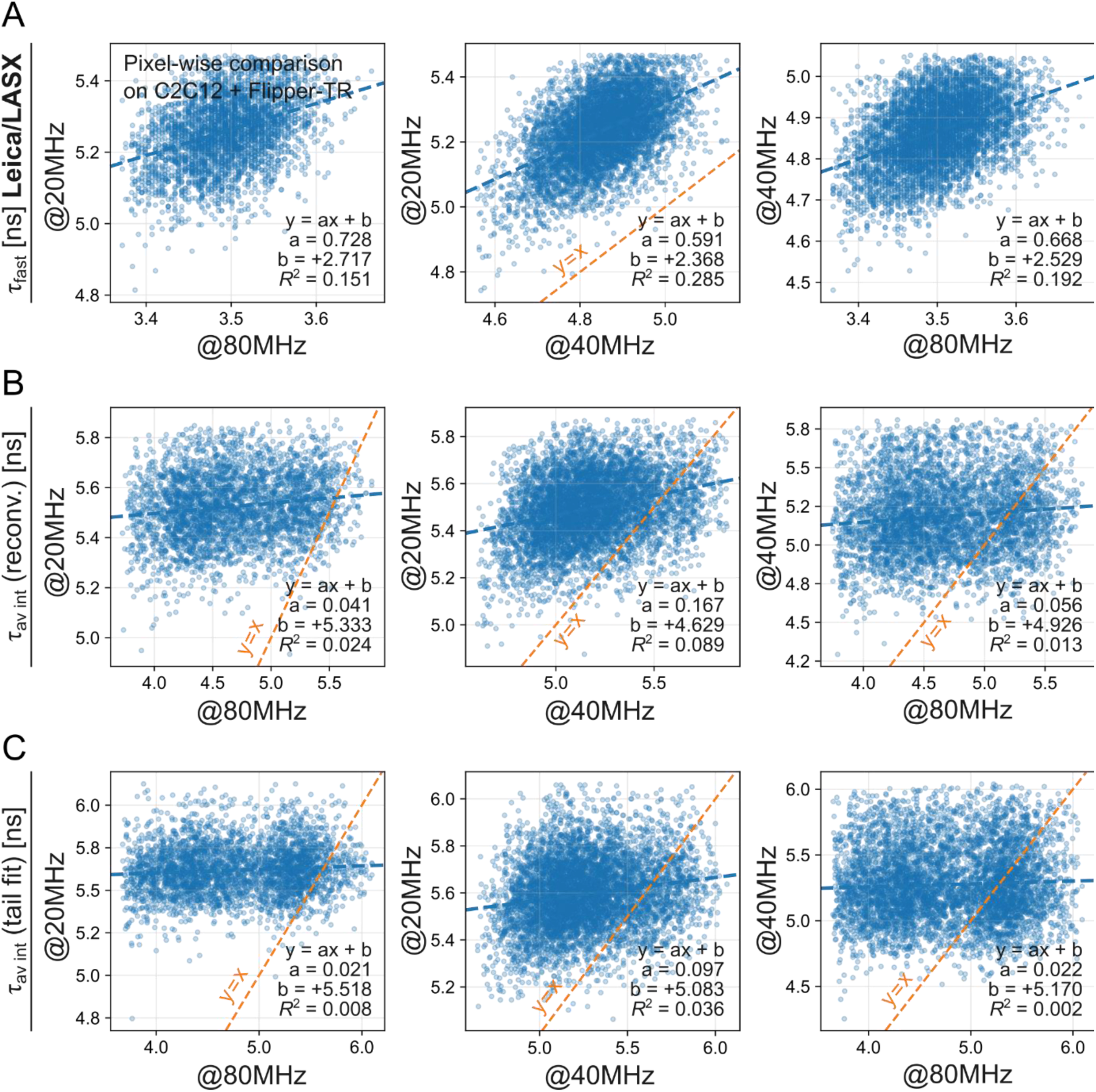
Correspondence between laser frequencies on Leica/LASX software. Rows (top to bottom): lifetime estimators τ_fast_, τ_av int_ (reconvolution), τ_av int_ (tail fit). Columns (left to right): pixel-wise scatter of 20 versus 80MHz; 20 versus 40MHz; 40 versus 80MHz. Each dot is one pixel. Axes report lifetime (ns) with different estimators depending on the row. Points are overlaid with a weighted Huber regression, a robust linear fit (blue dashed) and a y=x identity line (orange dashed). Each panel annotates slope, intercept, and R². Inclusion filters: lifetime ∈ [2, 10] ns in both images; intensity > 2500 counts in both images; symmetric percentile clipping on lifetimes (1st–99th) applied after the hard thresholds.

**Figure 13:**
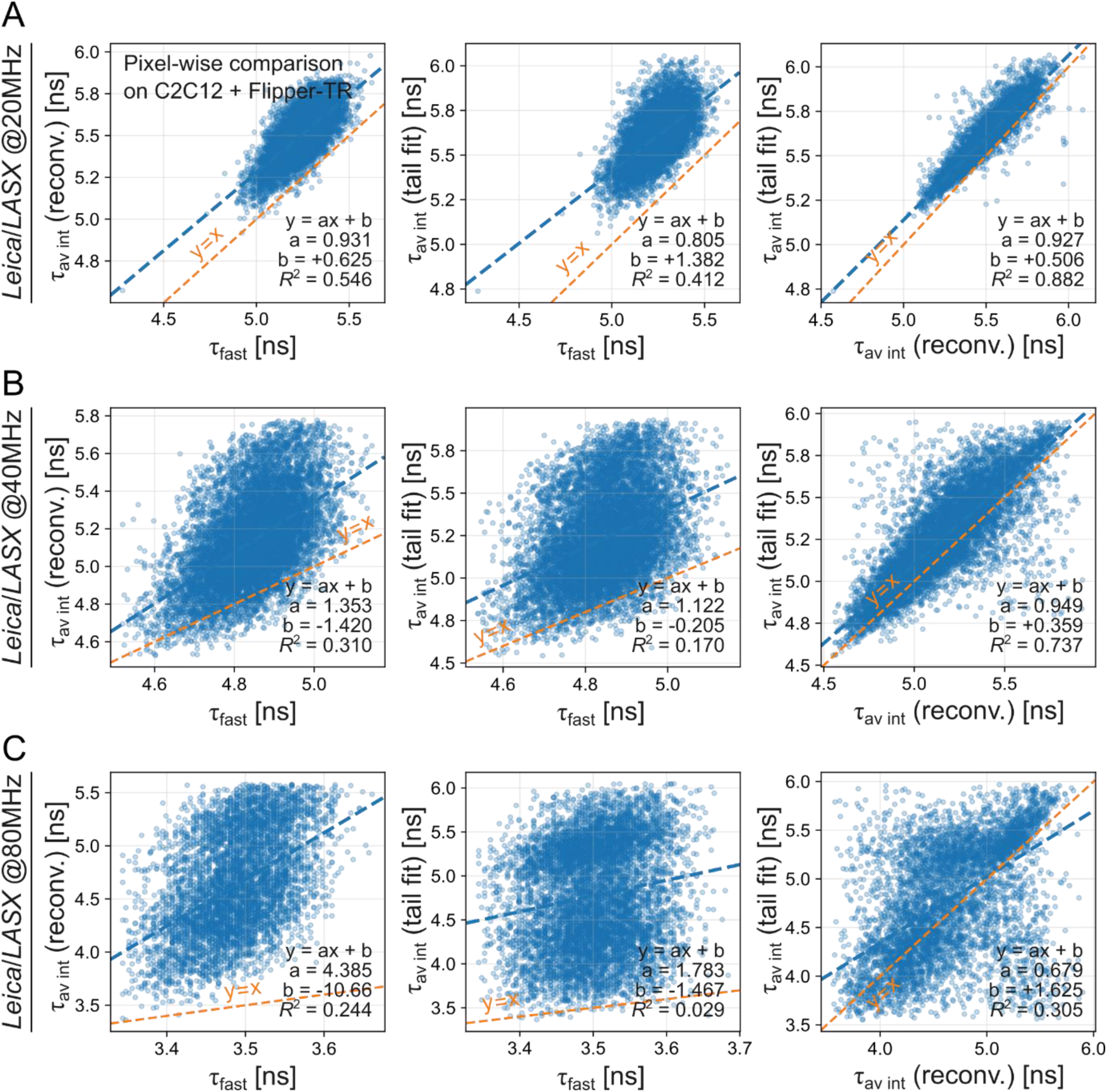
Correspondence between fit models on Leica/LASX software. Rows (top to bottom): laser repetition rates 20, 40, 80 MHz. Columns (left to right): pixel-wise scatter of τ_fast_ versus τ_av int_ (reconvolution); τ_fast_ versus τ_av int_ (tail fit) and τ_av int_ (reconvolution) versus τ_av int_ (tail fit). Each dot is one pixel. Axes report lifetime (ns) with different estimators as indicated on the axes. Points are overlaid with a weighted Huber regression, a robust linear fit (blue dashed) and a y=x identity line (orange dashed). Each panel annotates slope, intercept, and R². Inclusion filters: lifetime ∈ [2, 10] ns in both images; intensity > 2500 counts in both images; symmetric percentile clipping on lifetimes (1st–99th) applied after the hard thresholds.

### 2.3 Flipper-TR lifetime variability depends on photon count

So far, we have shown that lifetime estimator values and their variability depend on the fitting approach, microscope configuration, and laser repetition rate. While these comparisons emphasize instrument-related variability, an equally important factor is the inherent photon statistics of the measurement. Because TCSPC devices record one photon at a time, each photon carries only a small fraction of information about the underlying fluorescence decay, the number of detected photons directly constrains the precision of any of the lifetime estimators, i.e., at low photon counts, the variability of lifetime estimations such as fast lifetime is high, and values converge at higher photon counts (Figure 14). Generally, lifetime estimation inherently involves a trade-off between precision, photon count, and biological complexity. This is a classic problem with FLIM measurements (Esposito, 2020; Köllner & Wolfrum, 1992). In this section, we analyze numerically how photon counts influence the variability of the mean-arrival-time τ_fast_, the intensity-weighted average lifetime τ_av int_ from bi-exponential reconvolution, and τ_av int_ from bi-exponential tail in two different setups.

**Figure 14:**
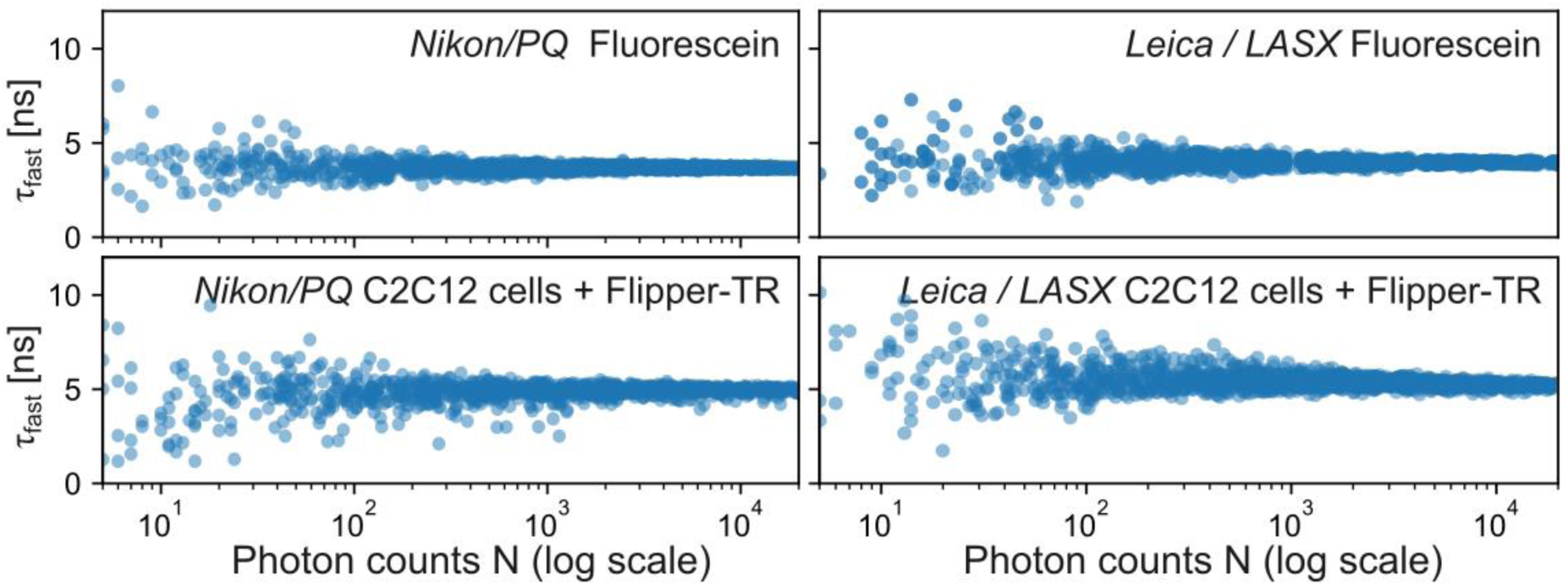
Fast lifetime values as a function of photon count. Per-pixel scatter of fast lifetime (τ_*fast*_, ns) versus photon counts for four datasets: fluorescein acquired on a Nikon–PicoQuant system (top left), fluorescein on Leica LAS X (top right), C2C12 cells stained with Flipper-TR on Nikon–PicoQuant (bottom left), and the same on Leica LAS X (bottom right). Each dot is one pixel. Pixels with low photon counts exhibit broad dispersion in τ_*fast*_, which narrows as photon counts increase

A given lifetime estimation is built from two stochastic layers: i) instrumental shot noise, and ii) biological heterogeneity. Let us consider a simple model to describe the variability of a lifetime estimate τ^_*ij*_ from replicate *i* in condition *j*. The lifetime estimate will depend on both the shot-noise α and the biological variability β, which are independent:

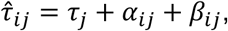

where τ_*j*_ is the true underlying lifetime value for condition *j*. In conditions with no biological variability, such as fluorescein standards or homogeneous samples, the only source of variability is shot noise. The precision of any unbiased fluorescence lifetime estimator is bounded by the Cramér–Rao lower bound (CRLB) (Kay, 1993, pp. 27–82), which states that the best possible precision (smallest possible standard deviation) of any unbiased estimator is inversely proportional to the square root of the information (in the case of FLIM, number of photons):

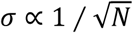

This relation is not a property of a specific FLIM estimator; it’s a general property of how random variables accumulate information. Nevertheless, the variability can still be influenced by the experimental and analytical context. For that reason, we include a photon-efficiency constant *c* that depends on system parameters and experimental design. The factor *c* is model-dependent and reflects how efficiently a particular fit extracts that information. The variance σ^2^ of τ^_*ij*_ can then be defined as:

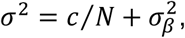

where *c* is the shot-noise constant, calculated empirically from the slope of σ^2^ versus 1/*N*; *N* is the photons collected for the given replicate τ^_*ij*_; and σ^2^ is the biological variance, which is zero for homogeneous standards such as buffered fluorescein solution.

This relation shows intuitively that more photons per record (higher *N*) will narrow the shot-noise term *c*Τ*N*, as shown in Figure 14. We should stop acquiring when *c* ∕ *N* ≪ σ^2^, as extra photons will add acquisition time and bleaching, but no statistical power. More replicates *n* will narrow the standard error of the mean by 1/√*n*. The optimal number of replicates depends on the statistical power and significance level needed. The best use of the “photon budget” will depend on the type of sample we are working with. In the case of homogeneous standards where the biological variability does not exist, the total photon budget *n* × *N* is what matters; we may trade photon counts *N* and experimental replicates *n* freely. In the case of biological samples, where biological variance σ^2^ is significant, we should first collect enough photons to make photon noise a small fraction of the total variance, then use the rest of the photon budget into extra biological replicates or larger spatial sampling.

We tested this model by acquiring series of fields of view acquired at increasing photon counts, and calculating for each acquisition the average photon counts and the variability of the pixel-by-pixel calculations of the mean-arrival-time τ_fast_, the intensity-weighted average lifetime τ_av int_ from bi-exponential reconvolution, and τ_av int_ from bi-exponential tail (Figure 15). To examine the influence of biological variability and microscopy setup, we acquired fluorescein standards and C2C12 cells stained with Flipper-TR at 20 MHz laser frequency rate in both Nikon/Picoquant and Leica/LASX setups (Figure 15A) at a pixel size of a few microns.

**Figure 15:**
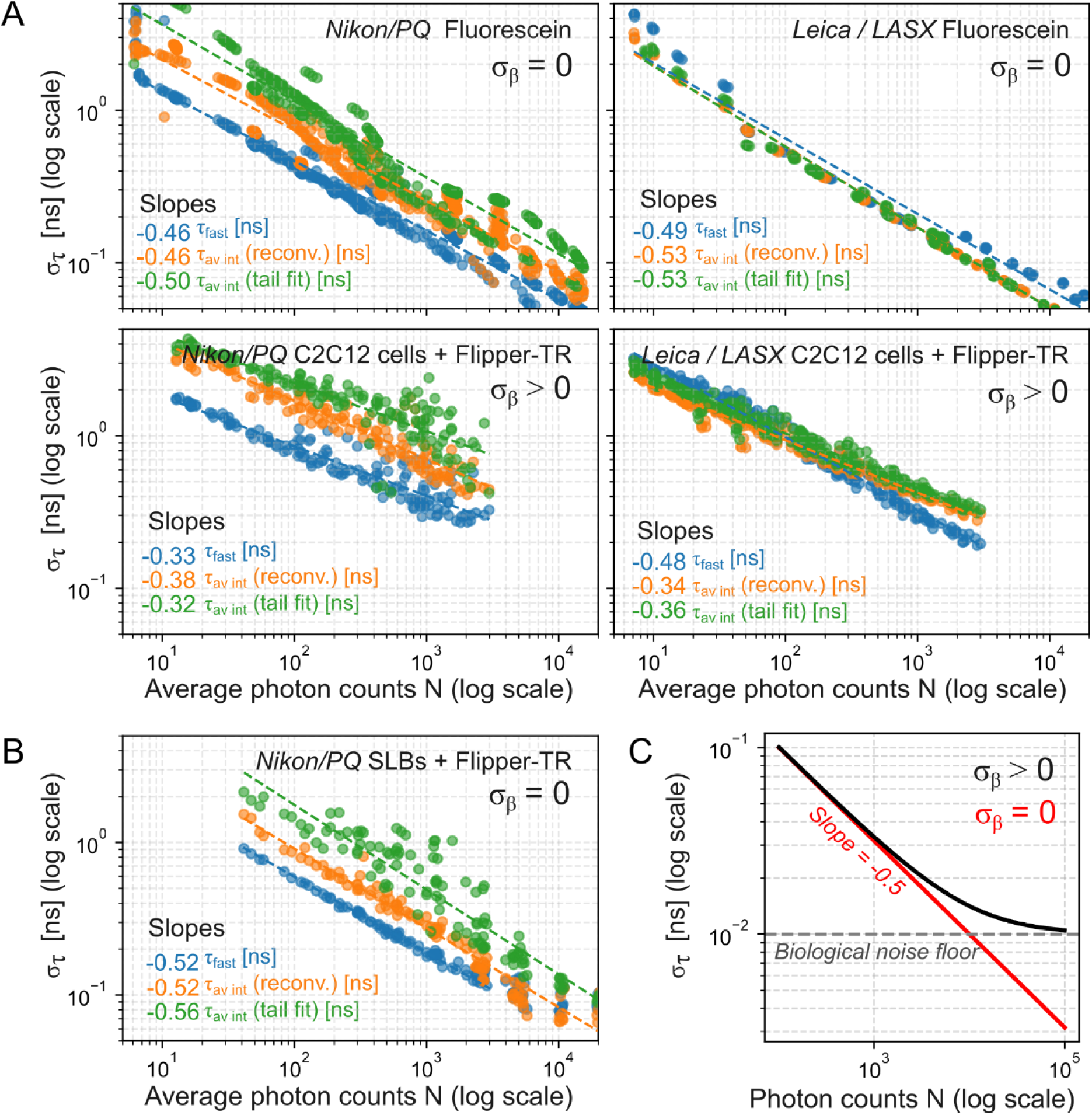
Lifetime estimator variability depends on photon count. (**A**) Per-field-of-view (FOV) scatter plots of lifetime variability versus photons for the same four conditions (top: fluorescein; bottom: C2C12 + Flipper-TR; left: Nikon–PicoQuant; right: Leica LAS X). The y-axis is σ_τ_ (standard deviation of lifetime across the image) and the x-axis is mean photon count per FOV. Points are colored by estimator: τ_*fast*_ (blue), intensity-weighted average lifetime from n-exponential reconvolution τ_av int_ (reconvolution) (orange), and the corresponding tail-fit τ_av int_ (tail fit) (green). Dashed lines show linear fits in log–log space; the slopes give the power-law exponent linking σ_β_ to average photon counts. Conditions having no biological noise (σ_β_≈ 0; fluorescein, top row) are indicated, whereas cells (bottom row) are expected to include biological noise (σ_β_ > 0). (**B**) Same analysis as in (B) for supported lipid bilayers (SLBs) made from DOPC with Flipper-TR. (**C**) Simple simulation of σ_τ_ versus photon counts (log–log). When biological noise is present (σ_τ_> 0; black), the curve deviates from pure shot-noise scaling and plateaus at a noise floor. When biological noise is absent (σ_τ_ = 0; red), σ_τ_ follows the shot-noise limit with a slope −0.5.

The standard deviation as a function of the average photon counts shows a clear power-law in agreement with our model. For fluorescein standards in both setups, the plots exhibited a log–log slope of approximately −0.5, which aligns with the theoretical expectation, σ = *c N*^−0.5^ (Figure 15A, top row). In Nikon/Picoquant setup, τ_fast_, τ_av int_ (reconvolution), and τ_av int_ (tail) clearly separate, indicating that the photon-efficiency constant *c* depends on the estimator: lowest for τ_fast_, intermediate for reconvolution fit, and highest for tail fit. In Leica/LASX setups, the three estimators converged on the same line, indicating no significant differences in their photon-efficiency constant. In other words, at a given photon count, estimators have different precision levels in Nikon/Picoquant, but they have an equivalent precision in Leica/LASX. Note that the fast lifetime on Nikon/Picoquant has lower shot noise constant (better precision) than any of the estimators in Leica/LASX; but the fits from Nikon/Picoquant significantly increase variability above that (Table 3).

**Table 3:** Shot noise constants for different fitting algorithms and manufacturers, in nanoseconds.

|  | Nikon / Picoquant |  |  | Leica / LASX |  |  |
| --- | --- | --- | --- | --- | --- | --- |
|  | Fast lifetime | Reconv. | Tail fit | Fast lifetime | Reconv. | Tail fit |
| Shot noise constant<br>$\sigma / \sqrt{N}$ | 3.10 | 10.05 | 23.02 | 6.37 | 6.62 | 6.59 |

We then performed an equivalent measurement in cells labeled with Flipper-TR. The maximum average photon counts had to be reduced to a few 10^3^ counts, to avoid cell damage. Nonetheless, the trends observed on the photon-efficiency constant were conserved, but the slopes deviated from −0.5, showing values around −0.35 (Figure 15A, bottom row). To determine whether this deviation was due to the decay characteristics of Flipper-TR or the presence of biological heterogeneity, we performed a control experiment using supported lipid bilayers (SLBs) composed of DOPC (Figure 15B). These SLBs are pure lipid bilayers composed of a single lipid type attached *in vitro* to a glass surface, presenting little or no spatial heterogeneity. The same analysis yielded a slope of −0.5, recovering the σ ∝ *c N*^−0.5^relation. This comparison shows that the deviation in cells originates from biological heterogeneity rather than from the probe itself. In cells, Flipper-TR samples a distribution of local membrane environments, which broadens the measured lifetime distribution and reduces the apparent slope. Thus, SLBs confirm both the theoretical power-law dependence and the added contribution of biological variability in cellular measurement (Figure 15C). Nevertheless, the slope of the data from cells does not visually plateau to a biological noise floor. This observation confirms that pixel-by-pixel image fits taken at reasonable phototoxicity (i.e. up to 1000s of counts per pixel) have still a significant impact from both the shot noise and the biological variability.

### 2.4 Practical application: analysis of membrane tension during osmotic shocks

As we can infer from the non-biological samples in Figure 15A, the standard deviation caused by the shot-noise takes small values (under 0.1 ns) only over 10^4^-10^5^ photon counts. In our experience, this photon count on live Flipper-TR samples is impossible to achieve at small pixel sizes without causing significant photodamage. One possible strategy is to work with bigger pixel sizes, perform region-of-interest estimations grouping several pixels, or even full-image lifetime estimations.

To put this in practice, we performed a 2-second interval time-lapse FLIM on HeLa cells undergoing a series of osmotic shocks (Figure 16-17). Before imaging, the cells were incubated 10 minutes with Flipper to ensure incorporation into the plasma membrane. Hypo-osmotic (∼300 to 150 mOsm by water addition) and hyper-osmotic shocks (∼150 to 533 mOsm by sucrose addition) were then sequentially applied during simultaneous FLIM acquisition.

**Figure 16:**
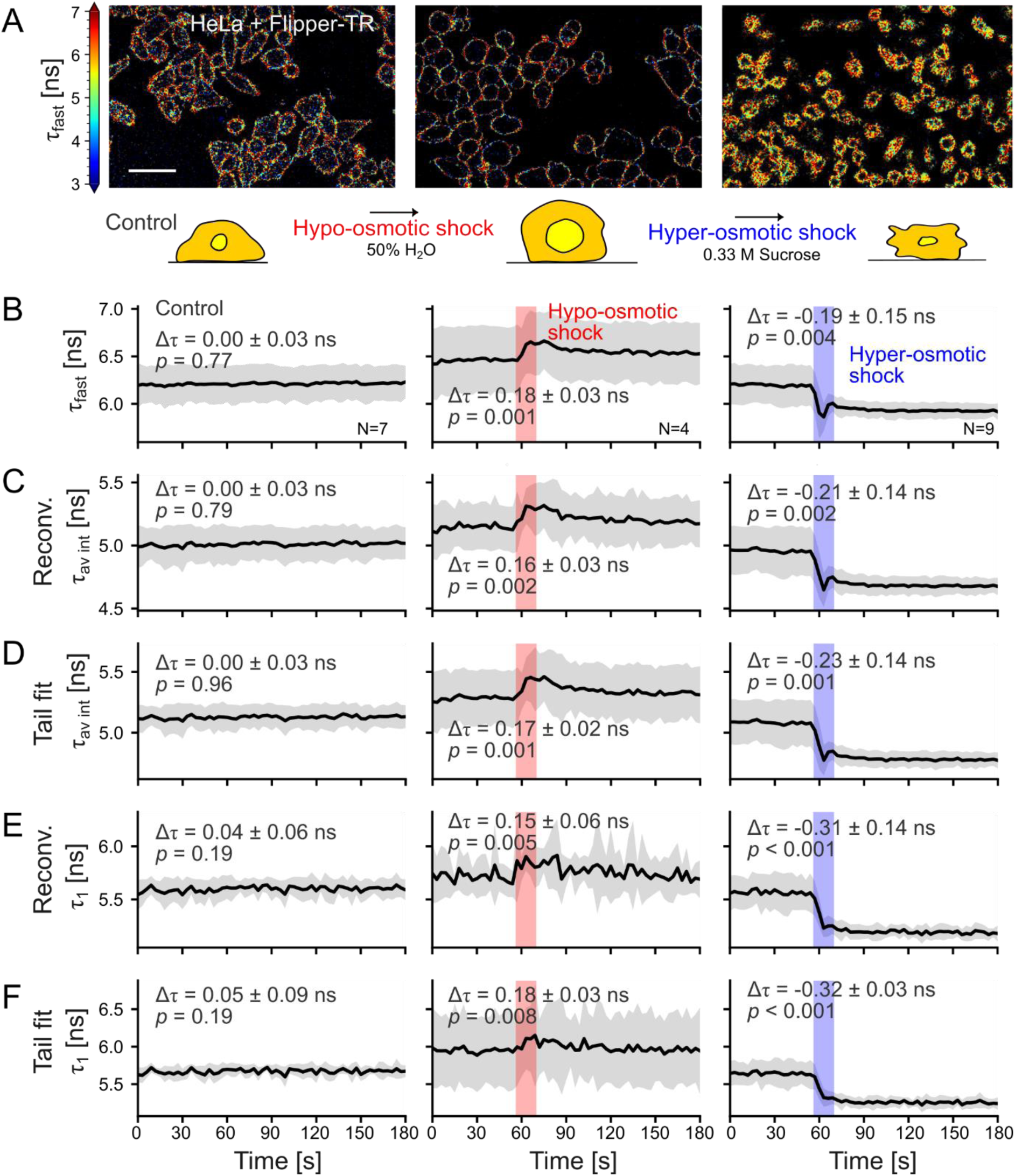
Comparison between analysis strategies of osmotic shocks. (**A**) Fast-lifetime (τ_*fast*_, ns) FLIM images of HeLa cells in three conditions: control (left), hypo-osmotic shock (50% H₂O; center), and hyper-osmotic shock (0.33 M sucrose; right). A shared color scale bar (ns) is shown on the left. Low-photon pixels are overlaid in black on the pseudo-colored τ_*fast*_ images. A schematic beneath the images summarizes the sequential protocol (control → hypo → hyper). (**B**–**F**) Time courses from the same recordings (3 min at 3 s per frame) shown as three subpanels per row: control (left), hypo (middle), hyper (right). The x-axis is time (s). The y-axis shows the across-replicate mean of the indicated lifetime estimator (ns); shaded bands indicate the inter-sample standard deviation. Whole-image values are used (no segmentation or masks). N denotes independent experiments. Estimators: (**B**) τ_*fast*_ (intensity-weighted mean arrival time); (**C**) intensity-weighted average lifetime τ_av int_ from a 2-exponent tail fit; (**D**) intensity-weighted average lifetime τ_av int_ from a 2-exponent reconvolution fit; (**E**) τ_1_ from a 2-exponent tail fit; (**F**) τ_1_ from a 2-exponent reconvolution fit. All lifetimes are reported in nanoseconds. Scale bar, 50 µm.

**Figure 17:**
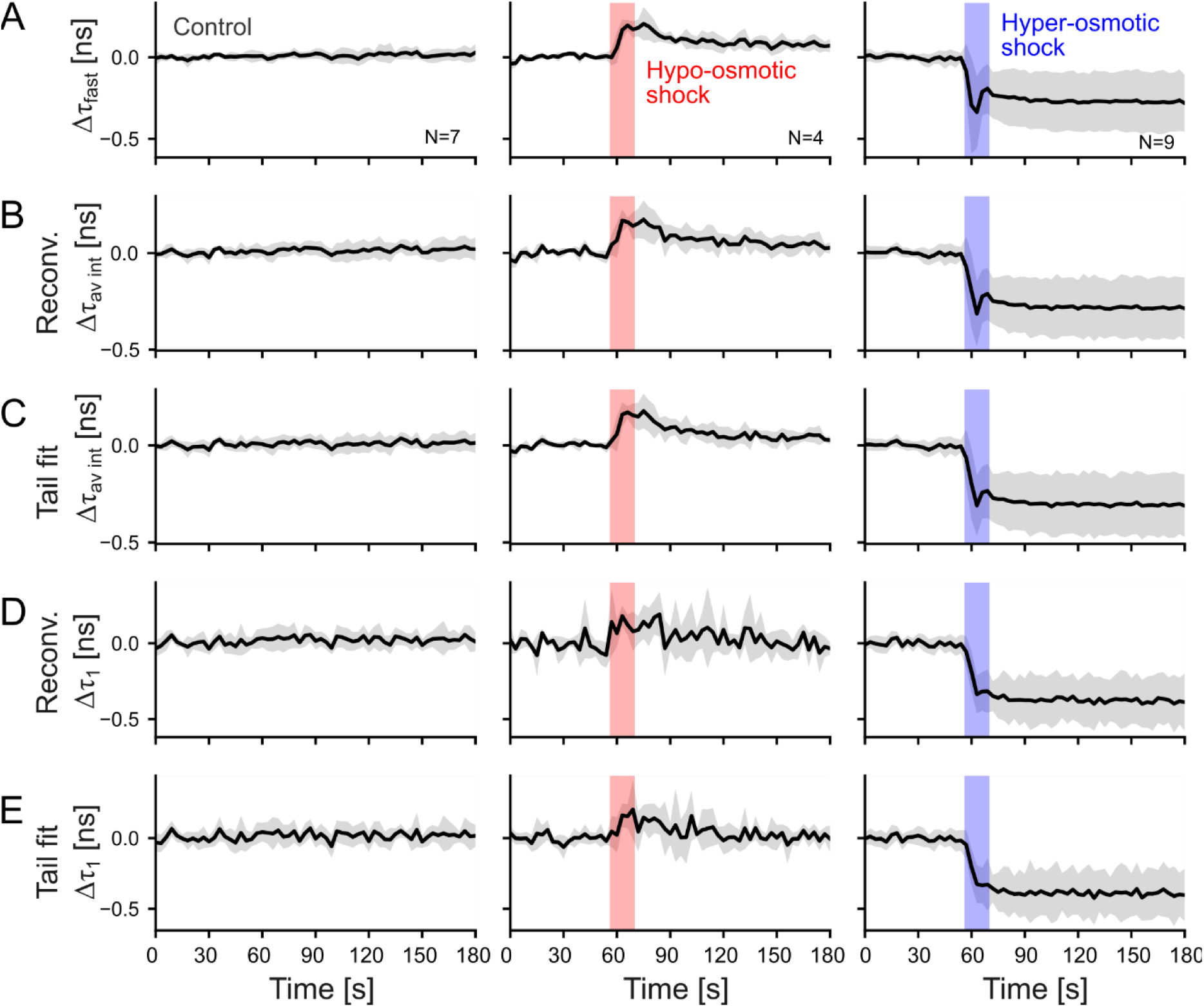
Lifetime offsets during osmotic shocks. Same as in figure 16 panels B-F, but values have been normalized. The mean lifetime from 0 to 45 seconds is calculated and subtracted from the entire time series. (**A**) Δτ_fast_ intensity-weighted mean arrival time; (**B**) Δτ_av int,_ intensity-weighted average lifetime from a 2-exponent tail fit; (**C**) Δτ_av int_, intensity-weighted average lifetime from a 2-exponent reconvolution fit; (**D**) Δτ_1_ from a 2-exponent tail fit; (**E**) Δτ_1_ from a 2-exponent reconvolution fit. All lifetimes are reported in nanoseconds.

In a previous study, it was reported that exposure of flipper probes to blue light can lead to the generation of reactive oxygen species (ROS). The accumulation of ROS promotes lipid hydroperoxidation within the membrane, which ultimately translates into changes in lifetime (Torra et al., 2024). Therefore, a control time-lapse with no perturbation was recorded to ensure no photodamage was caused (left panels on Figure 16). The photodamage and this time resolution imposes a very low average photon count of only 5 to 10 photons per pixel. Following again the curve from Figure 15A at this photon number, the standard deviation of the lifetime would be in the order of a few nanoseconds. This high noise is visible on the pixel-by-pixel fast lifetime calculations (Figure 16A). To remediate this, we can collect all photons to include them in a single estimation for the entire field of view. In this example, the 256×256-pixel image at 5 to 10 photons per pixel yields a few 10^5^ photons. This total photon count ensures that the shot-noise standard deviation is kept below 10^-2^ ns (derived from Figure 15), allowing for the measurement of subtle lifetime changes.

We tested the effect of osmotic shocks on the Flipper-TR lifetime on the mean-arrival-time τ_fast_ (Figure 16B), and several estimators from bi-exponential fits: the intensity-weighted average lifetime τ_av int_ from reconvolution (Figure 16C), τ_av int_ from tail fit (Figure 16D), the longest lifetime component τ_1_ from reconvolution (Figure 16E) and the τ_1_ from tail fit (Figure 16F). In terms of absolute values, estimators of the same type displayed similar values regardless of the fitting algorithm: τ_fast_ (∼6ns), τ_av int_ (∼5.1ns) and the τ_1_ (∼5.6ns). The response of HeLa cells to osmotic perturbations has been well described in the literature (Roffay et al., 2021; Venkova et al., 2022). Upon hypo-osmotic shock, HeLa cells swell transiently and undergo a regulatory volume decrease (RVD) within tens of seconds, restoring initial volume and tension values. Upon hyper-osmotic shock, HeLa cells shrink and decrease membrane tension and cannot to recover the initial values at a short timescale. The drops or increases in lifetime were similar among all the indicators and qualitatively recapitulate the expected behavior. The for the τ_1_ components, which displayed higher differences in hypertonic shocks. τ_av int_ estimators were less noisy than τ1 as expected from the data in Figure 7, while τ1 was less sensitive to the changes of focus occurring during the shock, displaying a cleaner curve. When comparing the distributions, the inter-sample heterogeneity observed in τ_fast_ is broader than that derived from fitting methods, yet the temporal fluctuations are comparatively smaller. Nevertheless, when performing paired statistical tests before and after the shocks, the effect of the perturbations is evident and statistically significant in all the estimators (see the p-values on each plot of Figure 16).

Despite the statistically significant changes in the paired t-tests, on the average plots, the standard deviation eclipses the lifetime change (Figure 16). This is due to the inter-sample variability (0.2-0.5ns) often being in the range of the changes in lifetime we might expect from physiological perturbations (0.1-0.3ns). For this reason, we often represent and calculate a relative Flipper-TR lifetime instead of absolute distributions of lifetime values. This normalization is calculated as offsets or different from the average or the initial state before the perturbation. In the case of osmotic shocks, we subtract the lifetime average of each repeat before the perturbation (0-45 seconds) from the entire measurement series. The resulting plots, therefore, represent lifetime offsets from the initial state (Figure 17). The conclusions or the statistical tests do not change qualitatively, but the visual representation is much clearer.

Finally, we applied phasor analysis available at Leica/LASX to the osmotic shock experiments to evaluate this fit-free approach under photon-limited conditions. In the phasor representation, each pixel is mapped to coordinates (x,y) = (G,S) in the form:

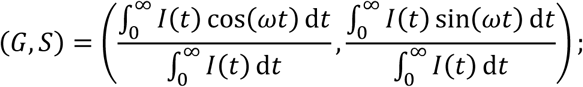

where *I*(*t*) is the fluorescence intensity decay function, i.e. the photon arrival histogram as a function of time after excitation; *t* is the time after each excitation pulse (on the nanosecond scale); ω is the angular modulation frequency of the laser excitation, ω = 2π*f*, where *f* is the laser repetition frequency (e.g., for 20 MHz:, ω = 2π ⋅ 20 × 10^6^ *Hz*; cos(ω*t*) and sin(ω*t*) are the basis functions of the Fourier transform at frequency ω (real and imaginary part, respectively); and ∫^∞^ *I*(*t*) d*t* is the normalization factor equal to the total number of detected photons. This formula is integrated into some commercial FLIM analysis software and provides a Fourier-space normalization of the decay histogram at the laser modulation frequency ω. The resulting phasor plots are 2D histograms that represent pixel frequencies as a function of the G,S coordinates. For mono-exponential decays, points fall exactly on the universal circle, with the arc position determined by τ, whereas mixtures of exponentials lie within the circle along line trajectories defined by linear combinations of their components.

In our phasor data, control, hypo-osmotic, and hyper-osmotic conditions yielded distinct shifts of the phasor clouds along the universal circle consistent with lifetime elongation upon swelling and shortening upon shrinking (Figure 18A). Temporal tracking of the (non-normalized) G and S coordinates (Figure 18B–C) qualitatively recapitulated the kinetics observed with fitted estimators, but without requiring model fitting and without normalization with a baseline. Notably, even at photon counts as low as ∼5–10 photons per pixel, the phasor distributions preserved directional shifts beyond the shot-noise dispersion. The high dispersion in the visual representation was reduced by an aggressive spatial and temporal binning (4 by 4 pixels and 3 time points).

**Figure 18:**
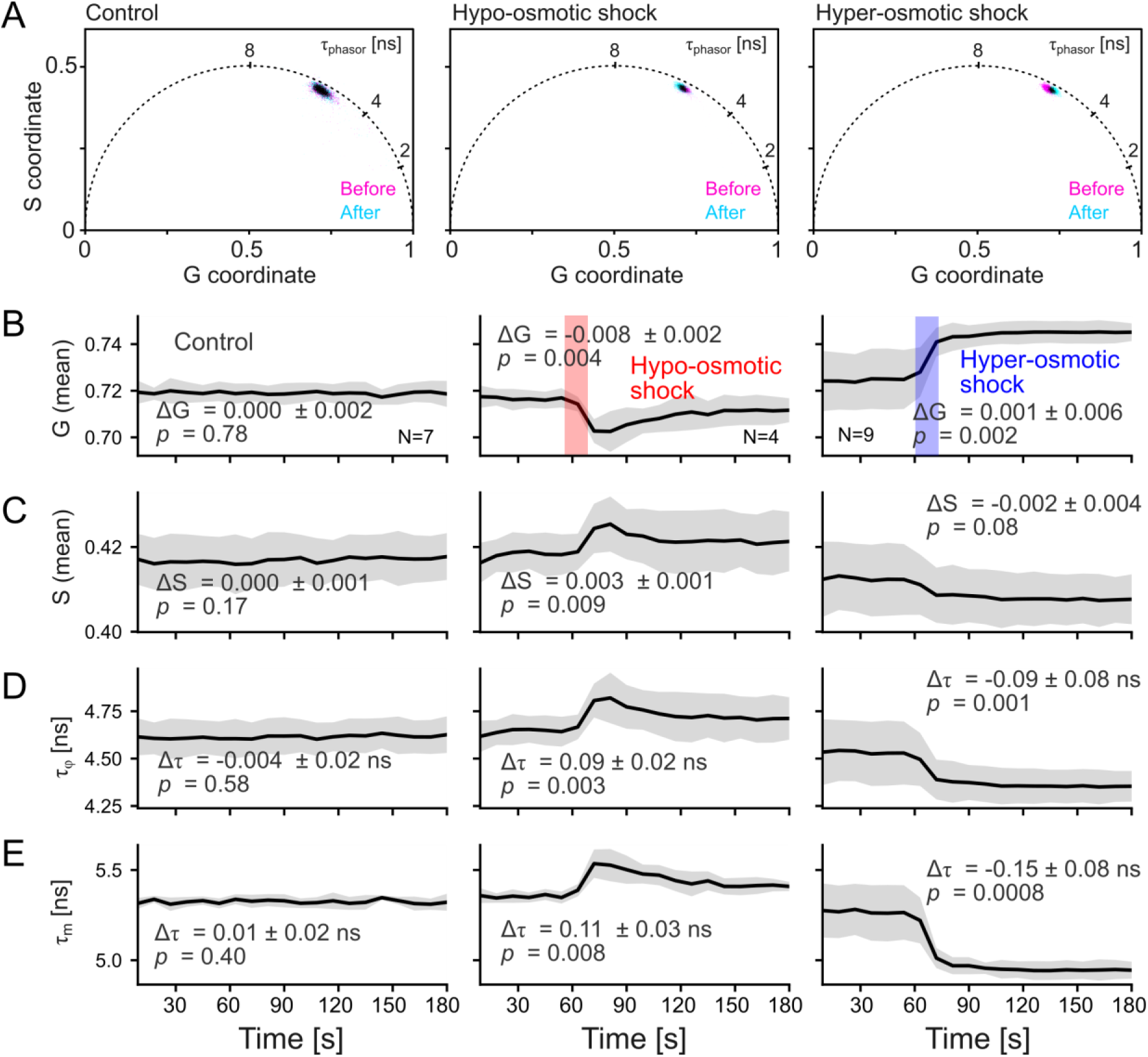
Phasor analysis of osmotic shocks. (**A**) Phasor plots before and after the sequential perturbation (as in Fig. 16): control (left), hypo-osmotic shock (50% H₂O; center), and hyper-osmotic shock (0.33 M sucrose; right). Pixel clouds are colored pink (before) and cyan (after); their overlap is shown in black. The universal circle is drawn as a dashed black line with lifetime markers at 2, 4, and 8 ns. Axes: G (x) and S (y). (**B**–**E**) Non-normalized time courses of the phasor coordinates from the same recordings, arranged as control (left), hypo (center), and hyper (right): (**B**) G versus time (s), (**C**) S versus time (s), (**D**) phase lifetime (ns), and (**E**) modulation lifetime (ns). Curves show the mean across biological replicates; shaded bands indicate inter-sample standard deviation.

We also calculated the first-harmonic phase lifetime τ_φ_ = *S*Τ*G*ω, where ω is the angular frequency (can be calculated from our laser frequency in MHz from 2π*f*_*MHz*_); and the first-harmonic modulation lifetime τ_*m*_ = √1 ∕ *m*^2^ − 1 ∕ ω, where *m* is the modulation depth (*m* = √*G*^2^ + *S*^2^). Both lifetimes are equal in the case of mono-exponentials, but in the case of Flipper-TR and it is multi-exponential both lifetimes display different values. Both lifetimes (Figure 18 D-E), but particularly the modulation lifetime display robust changes upon osmotic perturbation and remarkable consistency across samples as opposed to the fast lifetime and fit-derived values. The absolute varies in multi-exponents fits cannot be compared between frequency and time domain analysis, but they fall within similar range. Together, these results confirm that phasor analysis can serve as a reliable tool complementary to fitting, providing quantification of Flipper-TR lifetime changes during dynamic perturbations.

## 3. Discussion

### 3.1 Protocol to optimize acquisition and lifetime analysis

Based on the systematic benchmarking presented above, we propose below some guidelines for quantitative FLIM measurements and analysis of Flipper-TR across biological samples and instrument configurations.

#### Sample preparation

Samples are incubated with 50 nM Flipper-TR for at least 10 min in phenol-red–free medium. No wash step is required, as the probe is effectively non-fluorescent in aqueous solution. To monitor potential light-induced artifacts, a non-perturbed time-lapse control should always be recorded in parallel. Please refer to (Roffay et al., 2024) for further details on sample preparation, incubation, and Flipper-TR experimental design.

#### Acquisition settings

Excitation should be performed at 480–490 nm and emission collected at 575-625 nm. A repetition rate of 20 MHz (50 ns period) is optimal, as it accommodates the full decay of Flipper-TR. Higher rates (40–80 MHz) can bias long-lifetime components and should only be used for relative comparisons. In older setups sensitive to photon pile-up, photon count rates should be maintained below 10% of the repetition (more recent technologies such as rapidFLIM and FALCON lifetime hardware by Picoquant and Leica respectively are not likely to be affected by pile-up during Flipper-TR experiments). If absolute lifetime values are needed, we recommend performing sample calibration using a monoexponential standard such as fluorescein in PBS. For reconvolution fitting, a measured instrument response function (IRF) is also preferred to reduce errors: you can use a fluorescein solution quenched in saturated potassium iodide.

#### Photon budget and trade-offs

The precision of lifetime estimates is constrained by photon statistics, with standard deviation scaling as σ ∝ 1 ∕ √*N*. For pixel-wise mapping, a minimum of 1000 photons per pixel is recommended. Importantly, to reduce noise, a global fit can be used to define some parameter (e.g. the lifetime component values or only tau 2), and perform a pixel-wise fitting of the relative amplitudes to extract a pixel-wise average lifetime. For population-level comparisons or dynamic perturbations, aggregation of ≥10^5^ photons per field of view yields sub-0.01 ns precision without inducing photodamage. Optimal sampling therefore depends on whether spatial resolution or temporal resolution is prioritized.

#### Choose your lifetime estimator

*Fast lifetime* is simple to calculate, suitable for rapid quality control and analysis, but sensitive to repetition rate and artifacts, as all photons will be taken into account without any measure of quality control. *Bi-exponential fitting* yields the richest estimators. The intensity-weighted average lifetime provides the most stable and reproducible readout, whereas tau 1 (major component) is more variable but mechanistically interpretable. Reconvolution fits achieve higher accuracy with reliable IRF measurement, while tail fits are more robust when IRF estimation is absent. *Phasor analysis* is a model-free alternative particularly valuable at low photon counts or in experiments with complex decay patterns. The characteristics of each of the estimators we have benchmarked in this paper are summarized in Table 2.

#### Reporting standards

For reproducibility, Flipper-TR FLIM reports should specify probe concentration and incubation, microscope configuration (manufacturer, detector, repetition rate), photon budget per analysis unit, fitting model (tail vs. reconvolution, number of exponents), treatment of the IRF, estimator(s) reported, normalization procedure, and statistical tests. Cross-instrument comparisons of absolute values should be avoided unless supported by system-specific calibration.

### 3.2 Challenges and perspectives

Throughout this paper we have carefully benchmarked the analysis of Flipper-TR lifetime using realistic situations, including non-optimal laser acquisition, a variety of setups, low photon counts, or lack of calibration standards. Our benchmarking highlights several recurring challenges when performing quantitative FLIM with Flipper-TR, particularly under biologically realistic conditions. The first and most fundamental challenge is low photon counts. Because the probe localizes at the hydrophobic core of the bilayer and is sensitive to oxidation, excitation must be kept minimal to avoid photodamage (Torra et al., 2024). As a result, typical live-cell experiments yield only a few tens of photons per pixel, far below the photon numbers required for precise pixel-by-pixel fitting. In this regime, the standard deviation of lifetime estimates often overlaps with the physiological differences we aim to measure. As shown in Figure 15, there is no analytical workaround to overcome this limit; only by increasing photon counts through longer acquisition times, larger pixels, or pooling over regions of interest can precision be improved. Users should therefore adjust expectations: Flipper-TR FLIM experiments cannot be designed and interpreted as conventional confocal fluorescence imaging. Instead, lifetime analysis is more robust when performed on entire fields of view or large groups of pixels, or by averaging across many cells (García-Arcos et al., 2024).

A second challenge is variability across instruments and acquisition settings. Our data demonstrates that absolute lifetimes are not directly comparable across setups or laser repetition rates, even when using the same estimator. Whenever absolute lifetimes are of interest, calibration against mono-exponential standards (e.g. fluorescein in PBS) and a measured IRF are important. Otherwise, it is safer to report relative changes in lifetime (Δτ) normalized to an internal baseline (e.g. pre-perturbation period or control condition). This relative representation reduces instrument- and setup-dependent artifacts and makes results more reproducible across experiments.

A third challenge not directly addressed in this paper concerns data interpretation. As mentioned in the introduction, Flipper-TR is not a direct “membrane tension probe” but a lipid packing sensor. While lipid packing is strongly correlated with membrane tension in many contexts, it also integrates other physicochemical parameters including composition, phase separation, viscosity, substrate adhesion, and temperature. In practice, this means that lifetime shifts cannot be uniquely assigned to membrane tension without careful controls. Complementary experiments are required to disentangle the main driver of the changes in lipid packing, if such claims want to be made. These can be done by analyzing lipid composition, using Laurdan probes for polarity, molecular rotors for viscosity, or direct mechanical manipulations such as tether pulling when possible (Colom et al., 2018; Roffay et al., 2021). Importantly, lipid packing is itself a biologically relevant parameter and should not be dismissed even if the exact contribution of membrane tension cannot be fully resolved.

Beyond these technical considerations, it is important to recognize that our understanding of lipid packing dynamics in living systems remains incomplete. The behavior of lipid bilayers and of lipid-packing probes has been extensively characterized *in vitro* using well-defined compositions, where correlations with membrane mechanics are clear. However, in cells and tissues, lipid packing is regulated by a far more complex and dynamic interplay of tension, curvature, protein binding, trafficking, and metabolic state. How Flipper-TR and related probes respond to some subtle, physiologically relevant changes *in vivo* is not yet completely understood. This gap is especially evident in multicellular systems, where tension is coupled across membranes through adhesion, extracellular matrix contacts, and mechanical feedback between neighboring cells. The dynamic regulation of lipid packing in such contexts remains largely unexplored, and how packing-sensitive probes report on these processes is still an open question. Bridging this gap will require systematic integration of probe-based FLIM with mechanical perturbations, genetic manipulation, and quantitative theory.

In summary, the main challenges of Flipper-TR FLIM are low photon statistics, instrument-dependent variability, and interpretative complexity. These can be mitigated by prioritizing relative lifetime changes over absolute values, combining Flipper-TR with complementary probes, and adopting appropriate data analysis and visualization. At the same time, we emphasize that the biology of lipid packing regulation, particularly in multicellular contexts, remains a wide-open field, where the next steps will be not only technical but conceptual.

In the past years, Flipper-TR has moved from a specialist probe to a widely used reporter of lipid packing and membrane tension in live systems. This growth reflects also the growth of FLIM as a technique: broader access to FLIM platforms, clearer analysis choices, and better practices. By benchmarking estimators across instruments and repetition rates, and by providing practical controls and reporting standards, our study helps make Flipper-TR measurements more reproducible and comparable. Recent work is already using Flipper-TR to reveal rapid tension transients during cell shape changes, links between lipid order and mechanotransduction, and tension-dependent trafficking. We expect the workflows outlined here to support wider adoption in multicellular preparations and *in vivo*, where other tools to probe mechanical feedback are limited, and to help make Flipper-TR lifetime shifts a routine quantitative variable for biological discovery.

## Acknowledgements

The authors thank Jérémie Francfort (DIP Geneva Canton, Switzerland) for help and advice on the mathematical part of the work, and Naomi Sakai (University of Geneva, Switzerland), Mariano Gonzalez Pisfil (Ludwig Maximilian University of Munich, Germany), Raviv Dharan (University of Geneva, Switzerland), and Nelson Alonso Correa Rojas (EPFL, Switzerland) for critical reading of the manuscript and extensive feedback. The authors acknowledge funding from the European Research Council Synergy grant 951324-R2-TENSION.

## Author contributions

J.M.G.-A. conceived the study, co-acquired and exported the data, developed the analysis code, generated figure plots, and wrote the final draft of the manuscript. T.M. co-acquired and exported the data, produced the osmotic shock figures, and contributed to the draft version of the text. A.R. contributed through discussions of the project and provided critical feedback on the manuscript. All authors reviewed and approved the final version of the manuscript.

## Materials and methods

### Statement of the use of generative AI

The authors used Grammarly and ChatGPT (GPT-5, accessed September 2025) to assist with language editing, rephrasing of the main text and ChatGPT for improving the clarity and comments of the Python code. All scientific content and conclusions were produced by the authors.

### Resource availability

#### Lead contact

Further information and requests for resources, raw data, or analysis scripts should be directed to the corresponding author, Juan Manuel García-Arcos.

#### Materials availability

This study did not generate new reagents. Flipper-TR was commercially obtained from Spirochrome. Standard solutions (fluorescein sodium salt, sucrose, PBS) and lipid reagents were purchased from commercial suppliers (see below).

#### Data and code availability

All raw FLIM image files (`.nd2`, `.lif`, `.ptu`), fitted parameter outputs, and custom Python scripts used for post-processing and outputs are deposited in an open Figshare repository DOI: 10.6084/m9.figshare.30192961. Any other files are available upon request.

##### Cell culture and staining with Flipper-TR

C2C12 myoblasts (ATCC CRL-1772) and HeLa cells (gift from the lab of Matthieu Piel, Institut Curie, Paris) were cultured in DMEM supplemented with 10% FBS and 1% penicillin-streptomycin at 37°C, 5% CO2. For FLIM experiments, cells were plated onto 35-mm glass-bottom dishes (MatTek) and incubated for at least 10 min in phenol-red–free Leibovitz’s L-15 medium containing 1 µM Flipper-TR from a tock solution at 1mM as instructed by the manufacturer (Spirochrome #SC020). No washing was required, since Flipper-TR is non-fluorescent in aqueous solution. While imaging, samples were maintained at 37 °C using a stage-top incubation system.

For calibration experiments, fluorescein sodium salt (Sigma #F6377) was dissolved at 50 µM in PBS and imaged under identical conditions. Supported lipid bilayers (DOPC) and giant unilamellar vesicles (GUVs) were prepared as previously described (García-Arcos et al., 2024) and stained with 1 µM Flipper-TR to provide homogeneous reference samples.

##### Osmotic shock experiments

To probe lifetime sensitivity to membrane tension, HeLa cells stained with 1 µM Flipper-TR were imaged continuously during sequential osmotic perturbations. After baseline acquisition (0–45 s), a hypo-osmotic shock was applied at 60 seconds by diluting the medium 1:1 with sterile water (∼150 mOsm). After stabilization, a hyper-osmotic shock was induced by adding 1M sucrose solution to a final concentration of 0.33 M sucrose to reach ∼533 mOsm. A control time-lapse with no medium change was performed before the perturbations to assess phototoxicity. Images were acquired with ∼5–10 photons per pixel, and each perturbation was acquired in a different field of view of the same plate, i.e., one control, hypo-osmotic, and hyper-osmotic time lapses were acquired. For statistical analysis, lifetime traces were also normalized to the average baseline value and expressed as Δτ (see Figure 17).

#### Image acquisition and microscopy setups

##### Nikon Eclipse Ti2 – PicoQuant TCSPC

FLIM was performed on a Nikon Eclipse Ti2 inverted microscope equipped with a point-scanning A1 confocal system, Apo LWD λ 40×/1.15 WI objective (#MRD77410), and a perfect focus system. The setup included a stage-top OkoLab incubation chamber. Excitation was provided by a pulsed 485 nm diode laser (PicoQuant LDH-D-C-485) driven by a Sepia PDL-828, operated at 20, 40, or 80 MHz. Imaging parameters: pixel dwell 3.16 µs, scan speed ∼100 Hz, pinhole 1.2 Airy Units (AU). Emission was filtered through a 600/50 nm bandpass and detected with a PMA Hybrid 40 detector. Photon arrival times were recorded with a MultiHarp 150 TCSPC module. Control and acquisition were via NIS-Elements AR 3.30.05. No IRF measurement was used as input, but software generated

##### Leica Stellaris 8 FALCON

FLIM was also performed on a Leica Stellaris 8 FALCON microscope using a 40× water immersion objective. Excitation was provided by a white light laser tuned to 488 nm, and emission was collected from 500–600 nm with HyD detectors. Repetition frequencies of 20, 40, and 80 MHz were tested. Imaging parameters: frame repetition 8, pixel size 866.7 nm, pixel dwell 3.16 µs, scan speed 400 Hz, pinhole 1.2 Airy Units (AU). Samples were maintained at 37 °C in a stage-top chamber. Osmotic-shock experiments were conducted only on this system (Figures 16–18). No IRF measurement was used as input, but software generated

##### Leica SP8 DIVE – multiphoton

Two-photon FLIM imaging was performed on a Leica TCS SP8 DIVE upright multiphoton microscope using a HC FLUOTAR VISIR 25×/0.95 WI objective. Excitation was at 960 nm (fixed 80 MHz repetition rate), with emission collected by HyD-RLD2 detectors. Imaging parameters: frame repetition 8, pixel size 866.7 nm, pixel dwell 3.16 µs, scan speed 400 Hz, no pinhole on the external detectors. Microscope was maintained at 37 °C. No IRF measurement was used as input, but software generated

#### Step-by-step FLIM fit and export workflows

Below, we present brief workflows for data analysis and export in NIS/SymPhoTime, and LAS X. For more detailed guidance, please refer to the specialized and rich manuals and application notes provided by the manufacturers, but also methodological papers developed by third parties (Pandzic et al., 2022; Wang et al., 2024) or handbooks (Becker & Hickl GmbH, 2025).

##### Workflow A: NIS-Elements (Nikon) + SymPhoTime (PicoQuant)

Image acquisition was acquired in NIS-Elements AR 5.30.05. Analysis was performed in NIS-Elements AR Analysis 5.30.05 with the PicoQuant plug-in. Raw image data was saved as`.nd2`; recording the matching `.ptu` live during acquisition.

A1) Open and choose analysis route

1. Open the `.nd2` file in NIS-Elements Analysis.
2. Right-click and launch *PicoQuant plug-in* and choose one of:

- *Add an intensity channel* → creates a 2-layer stack (intensity + fast lifetime) suitable for quick inspection and export as `ome.tiff` (can be analyzed on Fiji, outside SymPhoTime).
- *Open in SymPhoTime* → launches PicoQuant SymPhoTime for full FLIM analysis and fitting options. A2) Initialize SymPhoTime FLIM
3. In SymPhoTime, the selected image/series appears in the workspace.
4. If analyzing a time series, select the frames to load.
5. Start FLIM via Imaging → *FLIM Start*.
6. Assess total photons (image-level) from the loaded dataset, displayed on the software. A3) ROIs, and binning
7. On the ROI tab, click new and define ROIs with any of the tools (*Freehand*, *Paint*, *Rectangle*, *Ellipse*, etc.). You can restrict analysis to selected ROIs or apply it to all.
8. Assess total photons (image-level) from your ROIs.
9. Adjust pixel/temporal *binning* as needed to optimize photon statistics for low-count regions (ideally keep settings consistent across comparable datasets). A4) Fast-lifetime (center-of-mass) maps
10. Run *Calculate FastFLIM* to compute fast lifetime maps.
11. Right click and *export* FLIM images *as OME-TIFF* for analysis outside the software. A5) Fitting analysis (Tail Fit / Reconvolution)
12. Choose n-Exponential Tail Fit or n-Exponential Reconvolution:

- *Set Model Parameters*: n = 2 for Flipper-TR.
- Set n = 1 for fluorescein.
- In this software, you do not pre-select τ components (τ_1_, τ_2_); the fit returns them.
13. Click *Initial Fit*, then *Fit All* to apply the model to all selected ROIs or frames. You can unclick certain components or constrain the range of parameters.
14. Always inspect χ² and residuals; if residuals show IRF-region structure, reconsider IRF handling or switch to Tail Fit.
15. After fitting, obtain the average lifetime (intensity-weighted average lifetime τ_*av int*_) or individual components from the results panel. A6) Exports, visualization, and quality control
16. Export fitted results (all parameters, or per-ROI if applicable) as ASCII for plotting and stats.
17. Visualize fitted and fast-FLIM images with customizable colour maps (e.g., Rainbow, RGB) and set colour bar ranges (min/max) consistently across samples.
18. Assess fit quality by inspecting:

- Chi-square (χ^2^), visible on the fit window but not exported along with fit data.
- Residuals (assess residuals structure versus noise).
- Lifetime histogram distributions.
- Fitted intensity decay curves (overlayed with fit).

#### Workflow B: LAS X FLIM-FCS Offline (Leica systems)

FLIM imaging was performed in LAS X software on the microscope, and offline analysis in LAS X FLIM-FCS Offline using .lif files.

B1) Open and organize datasets

1. Launch LAS X FLIM-FCS Offline and open the `.lif` file (Projects → Open).
2. If the dataset is a time series, select *All* and then *Split* in FLIM module to expose frames for per-frame analysis. B2) Fast FLIM (quick lifetime maps)
3. To analyze lifetimes rapidly, select the Fast FLIM module and compute τ_fast_ maps for each frame/FOV.
4. For cross-platform work, export raw images as OME-TIFF:

- To export select the individual range, enter 0.001 in the Lifetime section, and then saved the file in OME-TIFF format. The export will be 16-bit TIFF, therefore the factor 0.001 is important to have 3 decimals. 16-bit TIFF does not allow decimals. B3) Fitting (Reconvolution or Tail Fit)
5. Open Fit Model and choose either:

- n-Exponential Reconvolution (recommended when IRF is robust); enable IRF Background and IRF Shift to allow IRF fluctuations during fitting, or
- n-Exponential Tail Fit (robust when IRF estimation is uncertain).
6. Define the number of exponential components:

- n = 2 for Flipper-TR;
- n = 1 for fluorescein.
7. Click Fit, then Fit All to process the selected frames/ROIs.
8. Always inspect χ^2^ and residuals; if residuals show IRF-region structure, reconsider IRF handling or switch to Tail Fit.
9. Right click on results table and export all fit data to `.xlsx` B4) Pixel-resolved maps (FLIM Image Fit)
10. Go to FLIM Image Fit to generate per-pixel fit maps.
11. Adjust threshold and pixel binning to stabilize low-count pixels.
12. Allow fluctuations in τ_1_ and τ_2_ (enable those parameters). Importantly, to reduce noise, a global fit can be used to define some parameter (e.g. the lifetime component values or only tau 2), and perform a pixel-wise fitting with less free parameters.
13. Click Precise Fit to produce the final lifetime images. B5) Export results
14. After fitting, copy/export the fit table from Workspace (τ_1_, τ_2_, amplitudes, τ_av int_, χ^2^, errors) for plotting/statistics.
15. Export FLIM maps as OME-TIFF (or standard TIFF) for figure generation. B6) Phasor analysis
16. Open the Phasor module and choose Fast FLIM on the image visualization at the upper window.
17. Set time binning, pixel binning, and thresholds as needed.
18. To save for external analysis/plotting, right click on the fast FLIM image and select ‘Export raw image’.
19. Use *Save Phasor G S*. To export images select the individual range, entered 0.001 in the Lifetime section, and then saved the file in ImageJ compatible TIFF images.

- Note: in this software, only phasor images can be saved (not raw coordinate lists). If numerical G,S values are needed, extract them from the exported G/S images using our Python code or your own pipeline.

### Custom code for data analysis and figure generation

All custom analysis was implemented in Python (v3.11) as Jupyter notebooks. These notebooks, the input, and the output files are provided in the Figshare server (10.6084/m9.figshare.30192961) to ensure full reproducibility of the results presented here. Each notebook is organized around a specific figure or table in the manuscript, and is designed to process raw FLIM datasets exported from the acquisition software (SymPhoTime or LAS X) into either ASCII tables or OME-TIFF images. Unless otherwise stated, input files correspond to fluorescence lifetime image data (`.ptu` + `.nd2` for Nikon/PicoQuant, `.lif` for Leica), and outputs include processed lifetime values, statistical summaries, and publication-ready plots. Below we detail the purpose, inputs, outputs, and figure links for each notebook.

Figures_3_4_Plot_example_decays.ipynb

- Purpose: Demonstrates fitting of representative TCSPC decays (fluorescein and Flipper-TR) to mono- and bi-exponential models.
- Inputs: Photon arrival histograms and fits exported from SymPhoTime/LAS X as ASCII or OME-TIFF files.
- Outputs: Semi-log decay plots overlaid with fitted models, χ² residuals, and per-component lifetime values.
- Linked to: Figures 3–4 (example decay curves and fitting quality assessment).

Figure_6_tau_N.ipynb

- Purpose: Quantifies dependence of lifetime estimators (tau av int and tau 1) on number of exponentials used in the fitting model.
- Inputs: Exported fit data from SymPhoTime (fluorescein, Flipper-TR cells).
- Outputs: Scatter plots of tau av int and tau 1 versus exponential model complexity, with error bars.
- Linked to: Figure 6 (lifetime estimator stability across fit models).

Table_1_lifetime_averages.ipynb

- Purpose: Computes average lifetime values for Flipper-TR and fluorescein across instruments and fitting methods.
- Inputs: Exported lifetime tables from SymPhoTime and LAS X.
- Outputs: Aggregated summary table of tau fast, tau av int and tau 1 across conditions.
- Linked to: Table 1 (summary of lifetime averages across setups).

Figure_7_GUV_Fits_Correlation_Analysis.ipynb

- Purpose: Plots and tests correlation between tau av int, tau 1 and tau 2 in homogeneous GUV samples of different lipid compositions.
- Inputs: Bi-exponential fit tables exported from SymPhoTime for multiple GUV fields of view.
- Outputs: Linear regression plots of tau av int versus tau 1 and versus tau2, regression statistics (slope, intercept, R^2^).
- Linked to: Figure 7 (correlation analysis in GUVs).

Figures_8_9_FLIM_frequency_curves.ipynb

- Purpose: Assesses how lifetime estimators behave under different laser repetition rates (20, 40, 80 MHz).
- Inputs: TCSPC histograms for fluorescein and cells acquired at three repetition rates.
- Outputs: fast lifetime maps, decay histograms, fits, residuals plots.
- Linked to: Figures 8–9 (frequency dependence of lifetime estimators).

Figures_10_to_13_Frequency_Comparison_by_fit_by_brand.ipynb

- Purpose: Compares estimator behavior across frequencies and across microscope vendors (Nikon/PicoQuant vs Leica).
- Inputs: Pixel-wise lifetime maps (fast lifetime and tau av int from tail/reconvolution) from Nikon and Leica datasets.
- Outputs: Scatter plots with robust regression (Huber fit), slopes, intercepts, and R^2^ statistics.
- Linked to: Figures 10–13 (cross-frequency and cross-vendor comparisons).

Figure_14_15_std_plot.ipynb

- Purpose: Evaluates the scaling of lifetime variance with photon counts, testing against the Cramér–Rao bound.
- Inputs: Series of acquisitions at increasing photon counts (fluorescein, cells, supported lipid bilayers).
- Outputs: Log–log plots of σ_τ versus photon number N, fitted slopes, and overlay of theoretical scaling.
- Linked to: Figures 14–15 (precision analysis and variance modeling).

Figure_16_fast FLIM from tiff movie.ipynb

- Purpose: Extracts fast lifetime values from time-lapse FLIM of osmotic shock experiments.
- Inputs: OME-TIFF time series exported from LAS X (5–10 photons/pixel).
- Outputs: Whole-FOV fast lifetime time traces.
- Linked to: Figure 16B (fast FLIM time courses).

Figure_16_fit_analysis.ipynb

- Purpose: Plots osmotic shock time-lapse data.
- Inputs: Fit data from LAS X for each frame.
- Outputs: Time traces of different lifetime estimators after baseline subtraction.
- Linked to: Figures 16-18 (lifetime and phasor dynamics and normalized plots).

Figure_18_Phasor processing.ipynb

- Purpose: Processes phasor coordinates to extract average G/S per frame.
- Inputs: G/S phasor TIFF exports from LAS X.
- Outputs: Phasor plots with universal circle overlay, temporal trajectories of G and S.

Linked to: Figure 18 (phasor analysis of osmotic perturbations).

